# Brief high-fat diet exposure temporally reshapes affective behavior and prefrontal cortex function

**DOI:** 10.64898/2026.09.02.748924

**Authors:** Prince Peter Wormenor, Nathan C. Cupertino, Jonathan K. Crider, Jacob E. Turner, Whitney J. Walker, Brandon L. Roberts

**Author notes:** These authors contributed equally (Prince Peter Wormenor, Nathan C. Cupertino). **Correspondence should be sent to:** Brandon L. Roberts, Ph.D., University of Wyoming, 1000 E. University Ave, Biological Sciences, Room 316A, Laramie, WY 82071.

## Abstract

High-fat diet consumption is linked to a higher risk of obesity, diabetes, cardiometabolic diseases, psychiatric disorders, and cognitive dysfunction. Energy homeostasis and mood fluctuate throughout the 24-hour day, and perturbations in these rhythms have consequences for brain, body, and behavior. Our recent work demonstrates that layer 2/3 pyramidal neurons in the prelimbic prefrontal cortex (plPFC), a key region in affective behavior and cognitive function, undergo daily rhythms in their physiological function. In this study, we investigate how a one-week acute high-fat diet (aHFD) alters affective behavior and plPFC function. Further, we used time-of-day as a tool to determine how initiation of a HFD impacts the physiological properties of layer 2/3 plPFC pyramidal neurons in male and female mice. To explore these phenomena, we implemented the open-field test (OFT) and DeepLabCut to gain dynamic whole-body spatiotemporal resolution in our behavioral analysis. Next, we employed patch-clamp electrophysiology to determine how aHFD alters the intrinsic membrane properties and action potential dynamics of these neurons. Here we show that aHFD produces time-of-day dependent changes in exploratory behavior and physiology of plPFC pyramidal neurons, including altered intrinsic properties, firing profiles, and voltage-dependent membrane conductance in both sexes. These results lay the groundwork to explore the physiological mechanisms by which aHFD impacts plPFC function, affective behavior, and cognition. Overall, this study highlights the importance of understanding how diet impacts neural function before the onset of obesity.

## INTRODUCTION

Obesity and related metabolic disorders are among the most pressing global health challenges^1,2^. High-fat diet (HFD) consumption is a major contributor to obesity, type 2 diabetes, and cardiovascular disease, but its effects extend beyond cardiometabolic function^3^. HFD impairs cognitive function, disrupts emotional regulation, memory, and increases the risk of mood disorders^4–7^. Importantly, these changes can occur prior to significant weight gain, suggesting that the brain is an early and sensitive target of dietary perturbations. Understanding how acute exposure to HFD influences neural function is therefore essential for identifying early mechanisms linking diet to long-term health outcomes.

A growing body of work highlights the importance of circadian rhythms in regulating both energy balance and affective state^8–10^. Physiological and behavioral processes, such as feeding, locomotor activity, and stress responsivity, are regulated by daily rhythms, and their alignment with environmental light–dark cycles is critical for health^10–12^. Disruptions of these rhythms, whether by genetic, environmental, or dietary factors, exacerbate metabolic and psychiatric risks^9,13^. For example, circadian clock gene mutations are associated with hyperphagia, obesity, and altered mood-related behaviors^14–16^. While the impact of circadian disruption on metabolism is well-documented, far less is known about how metabolic challenges, such as short-term HFD exposure, affect time-of-day dependent changes in affective behavior and neurophysiology.

The prelimbic (pl) area of the prefrontal cortex (PFC) is a central hub for executive function, emotional regulation, and stress responses^17–20^. Dysfunction in this region is strongly linked to anxiety and depression^21,22^. plPFC pyramidal neurons exhibit diurnal rhythms in intrinsic excitability, suggesting that time of day influences how this region processes information and contributes to behavioral output^23,24^. Recent work further demonstrates cell-type and sex-dependent molecular rhythms in PFC pyramidal and parvalbumin interneurons, together with phase-dependent changes in PV-interneuron electrophysiology^25^. Perturbing these rhythms may alter cognitive and affective output, but whether short-term HFD disrupts diurnal PFC function remains poorly understood.

The PFC is also highly sensitive to dietary influences^21,26,27^. Prolonged HFD alters inhibitory neurotransmission, reduces GABAergic tone, and impairs executive function in other cortical regions^28^. In rodent models, even brief exposure to calorie-dense diets can alter hippocampal and cortical physiology, producing deficits in memory and impulse control^29–31^. However, most studies have focused on high-fat diet induced obesity. Much less is known about how short-term HFD affects the excitability and membrane properties of cortical neurons, or how these changes interact with daily rhythms in physiology. This represents a critical gap in understanding the earliest neural effects of dietary excess.

To address this, we investigated how brief HFD exposure impacts affective behavior and plPFC function in male and female mice. First, we employed high-resolution, machine learning–based analysis of the open-field test to quantify locomotion, exploration, and anxiety-like responses^32–34^. In addition to standard measures of distance traveled and zone occupancy, we implemented entropy-based metrics of speed and direction to capture variability and predictability of movement. These analyses allowed us to assess subtle changes in behavioral strategy that may be missed by conventional measures^35^.

We then combined these behavioral assays with patch-clamp electrophysiology in layer 2/3 pyramidal neurons of the plPFC, recorded at two distinct zeitgeber time (ZT) bins. This approach allowed us to test how aHFD modifies neuronal intrinsic properties, including resting membrane potential, resistance, capacitance, and action potential dynamics, in a time-of-day dependent manner. Given prior evidence for diurnal variation in PFC excitability^23,24^, this strategy provides a critical test of whether diet disrupts the rhythmic regulation of cortical neurons.

Together, this study addresses an important and underexplored question: how quickly does HFD exposure begin to impair cortical physiology and behavior? Our findings demonstrate that even short-term HFD produces time-of-day dependent changes in exploratory behavior and disrupts diurnal regulation of intrinsic and voltage-dependent properties in plPFC neurons. These results highlight that the brain is an early and sensitive target of dietary excess and underscore the importance of considering time-of-day interactions in studies of diet and brain function.

## METHODS

### Mice

All animal procedures for this study were performed in accordance with the National Institutes of Health’s Guidelines for the Care and Use of Laboratory Animals and with approval from the Institutional Animal Care and Use Committee (IACUC) at the University of Wyoming. Male and female wild-type mice on a C57BL/6J background (Jackson Labs, stock #000664) were obtained from our breeding colony at 8-12 weeks of age. Mice were randomly assigned to one of two experimental conditions in two different zeitgeber time (ZT) bins. All mice were group-housed prior to experiments in light-tight housing boxes at 25°C, under a 12:12-hr light:dark (LD) cycle and given food and water *ad libitum*. Prior to acute high-fat diet (aHFD) experiments, mice were singly housed in an LD cycle and then allowed to acclimate for one week before HFD exposure.

### Acute high-fat diet (aHFD)

Male and female mice were assigned to either standard control (CTR) chow (29%kcal protein, 58%kcal carbohydrate, 13%kcal fat; 3.36 total kcal/g) or a one-week acute (a) HFD (15%kcal protein, 26%kcal carbohydrate, 59%kcal fat; 5.49 total kcal/g; BioServ, Flemington, NJ) beginning at ZT0. Separate groups were used for behavior and electrophysiology experiments to avoid experiential confounds.

### Open-Field Test

The open-field test (OFT) was used to measure anxiety-like, locomotor, and associated behaviors. For our OFT design, we used standard dimensions of 50 cm x 50 cm x 38 cm (W x L x H)^33,36^. Matte white was chosen for consistency among studies and its non-reflective properties. The open-field and camera were calibrated to establish analysis zones and pixel to cm conversions prior to testing. The center zone was defined as the middle 40% of the arena floor. All mice were naïve to the OFT upon testing. They were placed in the center zone and allowed to explore freely for a short duration of 3 min. This duration was selected to capture the initial response to the environment and ensure that time-of-day is controlled while maintaining throughput for N values. An infrared camera connected to a Raspberry Pi was used for video acquisition. This prevented light exposure for tests occurring in the dark period.

### Behavioral Analysis

Behavioral assays were administered during the light (ZT3-6) or dark (ZT17-20) period. These time bins were selected to overlap with sacrifice and recording times of our electrophysiology experiments. Behavioral recordings were directly processed using DeepLabCut (DLC v3.0.0rc8) pose estimation outputs^34^. We utilized their SuperAnimal-TopViewMouse API implemented with a ResNet-50 pose estimation model and a Faster R-CNN MobileNetV3-Large FPN object detector^37^. The model tracks 27 body parts and was fine-tuned using video adaptation, with 4 adaptation epochs. This greatly increased prediction accuracy. For body-part specific analysis, we narrowed our analysis down to 5 distinct body parts: head midpoint, mouse center, mid-backend, tail base, and tail end. This encompassed movement for the entire length of the mouse and allowed for comparison between body parts and analysis of more nuanced changes in behavior. Data were filtered to only include predicted position of each part per frame with a prediction likelihood > 0.6. Tail segment five (tail5) was excluded due to unreliable tracking. Coordinates were converted from pixels to centimeters (cm) using a scaling factor derived from arena dimensions. For each frame, instantaneous displacement was computed as the Euclidean distance between consecutive coordinates. Arena zones were defined by experimenter determined boundaries (center vs. periphery), and each frame was assigned to a zone based on body part position. For radar plots, each variable was min-max normalized from 0 to 1 within the same sex and LD period across both diet groups; radar plots were used descriptively. Behavioral metrics were calculated as follows:

#### Locomotor Activity

Total distance traveled (cm) was calculated as the sum of all frame-to-frame displacements across the session. To control for changes in locomotor activity and any omitted frames (typically <10%), time spent in each zone was calculated as a percentage of total time in the testing environment. Average velocity (cm/s) was calculated as the mean instantaneous speed across all frames. Freezing was defined as movement < 1 cm/s. Freezing was quantified as the total duration of freezing episodes and expressed as a percentage of the total session time.

#### Speed Entropy

Instantaneous speed (cm/s) was calculated from frame-to-frame displacement multiplied by the frame rate. Speeds were grouped into 20 equally spaced bins spanning each mouse’s observed speed range. Shannon entropy^38^ was calculated as: 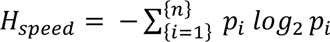 where *p*_*i*_ is the probability of observing speed in bin *i*. Higher entropy values indicate more variable locomotor output, whereas lower entropy indicates more stereotyped speed profiles^39,40^.

#### Directional Entropy

Movement direction at frame *t* was calculated as *θ*_*t*_ = *atan*2(*Δy*_*t*_, *Δx*_*t*_), preserving directional information across the full −*π* to *π* range. Shannon entropy (bits) was calculated as 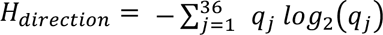 where *q*_*j*_ is the probability of movement in sector *j*. Higher entropy indicates greater variability in movement orientation^38–40^.

#### Avoidance Index

Zone occupancy was determined by frame-wise classification, and dwell time was calculated from the number of frames in each zone. The study-defined OFT avoidance index was: 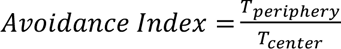 where *T_periphery_* and *T_center_* are dwell times in the respective zones; no analyzed mouse had zero center dwell. This index provides a composite measure of spatial avoidance by capturing the relative preference for peripheral versus center regions of the arena, with higher values reflecting greater center avoidance. For statistics, heatmaps, PCA, and behavioral radar plots, the avoidance index data was natural-log (ln) transformed to account for the large distribution of values from mice with a very low *T*_*center*_. Raw values are plotted and presented in tables. Radar plots were used descriptively.

### Brain slice electrophysiology

Mice from experimental groups (CTR and aHFD) were euthanized 1 hr before their respective ZT bin (i.e., mice were euthanized at ZT5 for recording bin ZT6-10), and brain slices were prepared as previously described^23,24^. For collections occurring during the dark period, mice were anesthetized in a dark box and quickly transferred to a dissection tray for euthanasia, which occurred in less than 60 sec. After euthanasia, the brain was immediately removed and the forebrain was blocked while bathing in a 0-4°C oxygenated N-methyl-D-glucamine (NMDG) cutting solution composed of (mM): 92 NMDG, 2.5 KCl, 1.25 NaH_2_PO_4_, 30 NaHCO_3_, 3 sodium pyruvate, 2 thiourea, 20 HEPES, 10 MgSO_4_, 0.5 CaCl_2_, 25 glucose, 20 sucrose. The cutting solution was brought to pH 7.4 with ∼ 17 mL of 5 M HCl^41^. The forebrain was then mounted and sectioned on a vibratome (VT1200S, Leica Biosciences, Buffalo Grove, IL, USA) with a sapphire knife (Delaware Diamond Knives, Wilmington, DE, USA) yielding 2-3 slices containing the plPFC from each mouse (250 μm). Slices were transferred and allowed to recover for 30-45 min in room temperature recording artificial cerebrospinal fluid (aCSF) solution composed of (mM): 124 NaCl, 3.7 KCl, 2.6 NaH_2_PO_4_, 26 NaHCO_3_, 2 CaCl_2_, 2 MgSO_4_, 10 glucose. aCSF had a final pH of 7.3–7.4, osmolarity of 307–310 mOsm, and was continuously bubbled using 95% O_2_/5% CO_2_. For recordings, brain slices were transferred to a perfusion chamber containing aCSF maintained at 34–37 °C with a flow rate of 1 mL/min. Neurons were visualized using an Olympus BX51W upright microscope (Evident Scientific, Tokyo, Japan). Recording electrodes were back-filled with an internal solution composed of (mM): 125 K-gluconate, 2 KCl, 5 HEPES, 10 EGTA, 5 NaATP, and 0.25 NaGTP. All internal solutions were brought to pH 7.3 using KOH at 297–301 mOsm. The calculated liquid-junction potential (LJP) was ∼15.9 mV, and all values in the manuscript are the raw values without the calculated LJP correction. Patch electrodes with a resistance of 3-8 MΩ were guided to neurons with an MPC-200-ROE controller and MP285 mechanical manipulator (Sutter Instruments, Novato, CA, USA). Patch-clamp recordings were collected through a MultiClamp 700B digital amplifier and Clampex recording software (Molecular Devices, San Jose, CA, USA). Current-clamp current-step protocols were performed from the cell’s endogenous resting membrane potential and used 1 sec 10-pA steps from −100 to +190 pA. Voltage-clamp voltage-step protocols were performed from a holding potential of V_H_ = −70 mV and used 10-mV steps from −120 to +30 mV. All compounds were obtained from Tocris, Cayman Chemical, Sigma-Aldrich, and Thomas Scientific.

### Statistical Analysis

#### Behavioral Measures

Behavioral measures derived from mouse-center tracking were analyzed separately by sex using diet × light/dark-period two-way ANOVA. Body-part-specific measures were analyzed separately by sex and period using mixed-model ANOVA, with diet as the between-subject factor, body part as the within-subject factor, and mouse as the subject. Greenhouse-Geisser-corrected p values were used for body-part specific effects when sphericity was violated. Sidak-corrected simple-effects comparisons were used for post hoc analysis.

#### Principal Component Analysis (PCA)

Behavioral and electrophysiological PCA were performed separately by sex using scikit-learn^42^. Variables were z-standardized before PCA^43,44^; no missing values were present in the behavioral feature matrix. The behavioral PCA included center-entry count, avoidance index, directional entropy, percent freezing, velocity, and speed entropy. The electrophysiological PCA included resting membrane potential (RMP), membrane capacitance (Cm), membrane resistance (Rm), action potential threshold, peak amplitude, afterhyperpolarization (AHP), half-width, rise tau, decay tau, action potential area, and rheobase. Diet and ZT were excluded from the feature matrices and used only to label score plots. Scores, variance explained, and loadings were reported for PC1 and PC2.

#### Chronometabolic Risk Index (CRI)

A study-specific exploratory^44^ CRI was calculated separately by sex from the same six behavioral features. Within each ZT bin, each feature was standardized to the same-sex CTR mean and SD, *z*_*ikt*_ = (*x*_*ikt*_ − *μ*_*CTR*,*kt*_)/*σ*_*CTR*,*kt*_. Feature signs (*s*_*k*_) were aligned so that the observed overall aHFD versus CTR deviation was positive, and the six standardized deviations were combined using normalized PCA-loading weights derived from the magnitude of each feature’s loadings across PC1-PC5; and 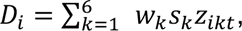 where normalized *w*_*k*_ comes from the magnitude of that feature’s PC1-PC5 loadings. The composite was then re-standardized across all animals of that sex, *CRI*_*i*_ = (*D*_*i*_ − *μ*_*D*_)/*σ*_*D*_. Because orientation used the observed diet contrast, the CRI was treated as a descriptive, outcome-oriented composite rather than a confirmatory measure. CRI scores were analyzed separately by sex using diet x time two-way ANOVA followed by Sidak-correct simple-effects comparisons within each time bin. Cohen’s d was calculated within each time bin to describe effect magnitude^45^.

#### Electrophysiology

Only neurons with input resistance >50 MΩ were included. Neurons were not considered for further analysis if series resistance exceeded 60 MΩ or drifted >10% during baseline. Rheobase was calculated as the first current step that elicited an action potential, and action potential properties were measured from the first evoked action potential to avoid variance from dynamics of repeated firing. Steady-state current (pA) from voltage steps (−120 to +30 mV in 10-mV increments; holding potential, −70 mV) was normalized to whole-cell capacitance (pF) to determine current density (pA/pF). Apparent reversal potential for current density was obtained by linear interpolation between adjacent voltage steps bracketing zero current; recordings without a single measured crossing were not extrapolated. Apparent slope-conductance density (nS/pF) was calculated over −120 to −90 mV and 0 to +30 mV and designated hyperpolarized- and depolarized-range conductance, respectively. Sexes were analyzed separately by diet × ZT two-way ANOVA with Sidak-corrected simple-effects comparisons.

Power calculations used G∗Power 3.0 software (Franz Faul, Uni Kiel, Germany) for a p-value of <.05 with 80% power, and expected effects from related experiments^23,24,46^. Adequate sample sizes were based upon expected effect sizes from similar experiments. Primary electrophysiology analyses treated cells as the analysis unit and figure symbols reflect these analyses. Animal numbers (N) are reported in the tables; values are presented as mean ± SEM. Fisher’s exact tests compared responsive-cell proportions. Analyses were performed in Python 3.11.3 using pingouin 0.6.1, statsmodels 0.15.0, seaborn 0.13.2, and matplotlib 3.11.

## RESULTS

### A one-week acute HFD alters time-of-day dependent behavior in male and female mice

Obesity contributes to psychopathologies, including anxiety in both humans and mice^4,5,47,48^. However, there are limited studies on how acute (a) HFD exposure impacts behavior^5,49^, and it is unknown how short-term HFD exposure impacts the diurnal properties of affective behavior. To test the impact of aHFD on exploratory behaviors, mice were placed on control (CTR) or aHFD and assigned to light (inactive) or dark (active) period prior to behavioral testing in the open-field test (OFT) (**Fig. 1a,b,e**). In male mice, center-zone entry counts, time spent in center zone, and total distance traveled were decreased during the dark period independent of diet treatment (**Fig. 1c; Tables S1,S2**). Further, male aHFD mice spent less time in the center zone, and this effect was restricted to the light period, resulting in a main effect of diet for the avoidance index (**Fig. 1c,d**). Male aHFD mice also showed an overall diurnal shift in velocity and decreased freezing behavior (**Fig. 1d**).

**Figure 1.**
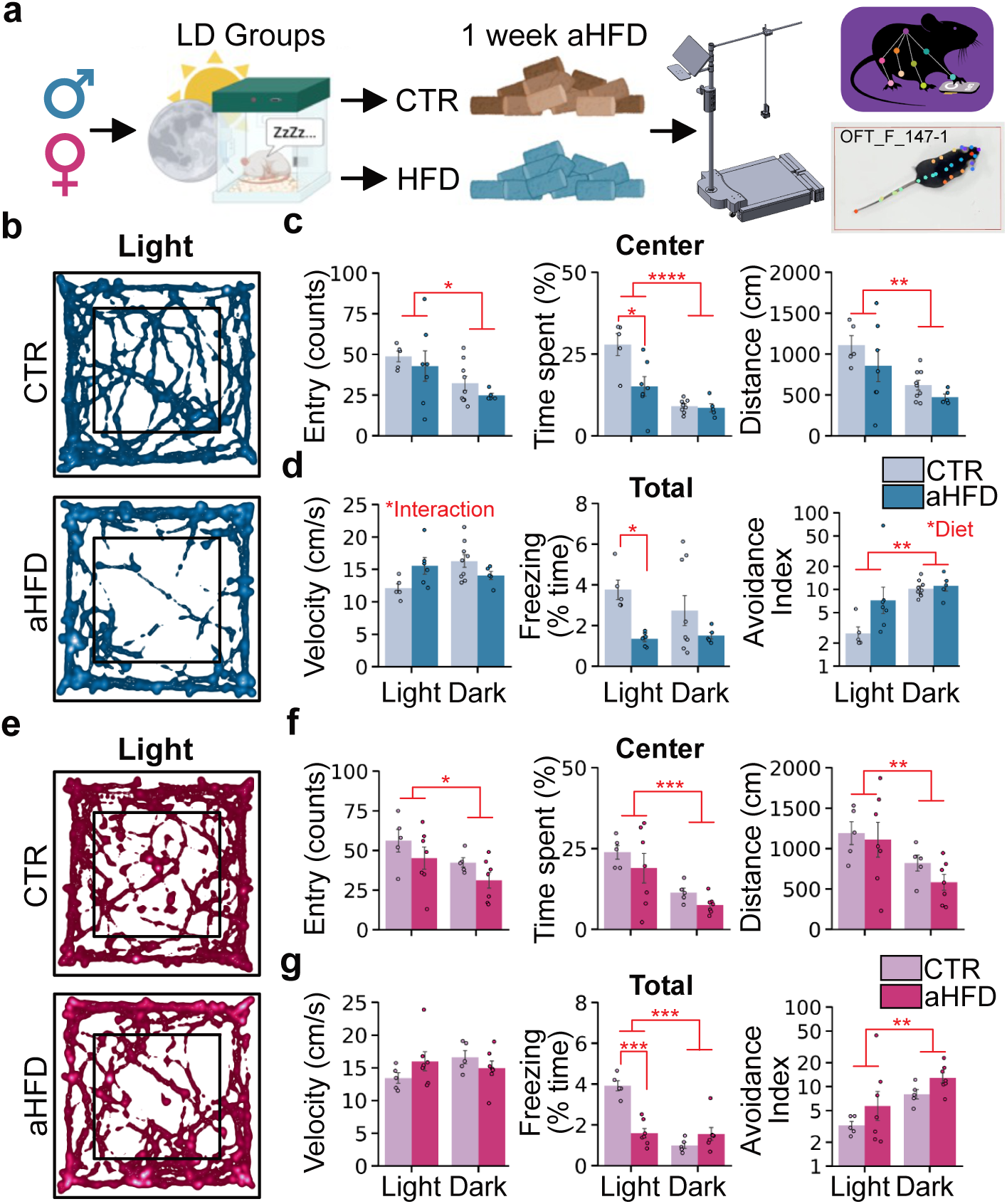
aHFD alters OFT performance in male and female mice. **(a)** Diagram of aHFD protocol for behavioral testing and video processing. **(b)** Single animal heatmap tracking of mouse-center during an OFT trial from a CTR (*top*) and aHFD (*bottom*) male and **(e)** female mouse. **(c)** Center zone entry counts (*left*), time spent (*middle*) and distance traveled (*right)* in CTR and aHFD male and **(f)** female mice during the light and dark period. **(d)** Total average velocity (*left*), freezing time (*middle*) and avoidance index (raw values are plotted on a logarithmic axis, and ln-transformed for analysis; *right)* in male and **(g)** female mice. *Two-way ANOVA for main effects of diet, ZT bin, and interaction **(Tables S1, S2)**. *p < 0.05, **p < 0.01, ***p < 0.001*.

Biological sex is a major factor in understanding how environmental perturbations impact physiology and behavior^23,24,50,51^. Thus, to address sex as a biological variable, we repeated these experiments using female mice (**Fig. 1e-g**). In female mice, there was an overall decrease in center entries, time spent in the center zone, and distance traveled in the center zone during the dark period independent of diet (**Fig. 1f; Tables S1,S2**). Overall freezing time was also greatly reduced in aHFD female mice during the light period (**Fig. 1g**). There was no main diet-effect for the avoidance index, as the LD period had the majority of main effects except for the diet effect of total freezing time (**Fig. 1f,g**).

### aHFD shifts overall behavioral phenotype in male mice

To address the complex behavioral interactions in movement between diet and light period in male mice, we extended our analysis to consider whole-animal movement across 26 body parts and split our analysis to focus on the light and dark periods separately (**Fig. 2a,g**). Qualitative observations from experimenters noted that aHFD mice appeared highly sporadic. To measure this observation, we calculated directional, speed, and overall entropy across five (5) body parts that capture the main segments of body movement (**Fig. 2b-d**). In the light period, there was a main effect of diet and body part for speed entropy and overall entropy (**Fig. 2b-d; Table S3**). Given these changes, we analyzed the avoidance index to account for whole-animal movement and behavior. When controlling for within-subject repeated measures (i.e., body parts), the avoidance index was not altered in aHFD mice during the light period, but displayed a strong trend (**Fig. 2e; Table S3**). We next plotted our primary metrics in a radar plot to visually observe overall behavioral shifts by normalizing the values in **Table S1** (**Fig. 2f**). Interestingly, when repeating this analysis for the dark period, there was no main effect of diet for entropy measures or avoidance index, resulting in a visually modest overall behavioral shift (**Fig. 2h-l**). To capture the overall structure of behavioral variation across multiple measures simultaneously, we performed a principal component analysis (PCA) that showed clear separation between CTR and aHFD during the light phase with PC1 (35.9%) and PC2 (28.4%) accounting for 64.3% of total variance (**Fig. 2m**). Of the top six loadings, PC1 was driven primarily by freezing time, speed entropy, and directional entropy, while PC2 was driven primarily by entry counts, avoidance index, and directional entropy (**Fig. 2n; Table S5**). To evaluate how time-of-day impacts these overall behavioral shifts, we calculated the exploratory chronometabolic risk index (CRI) as a PCA-loading weighted, directionally aligned composite of ZT-standardized behavioral measures. The composite showed its clearest separation between CTR and aHFD males during the light period (**Fig. 2o**), suggesting that aHFD-induced behavioral alterations were most pronounced during the light period.

**Figure 2.**
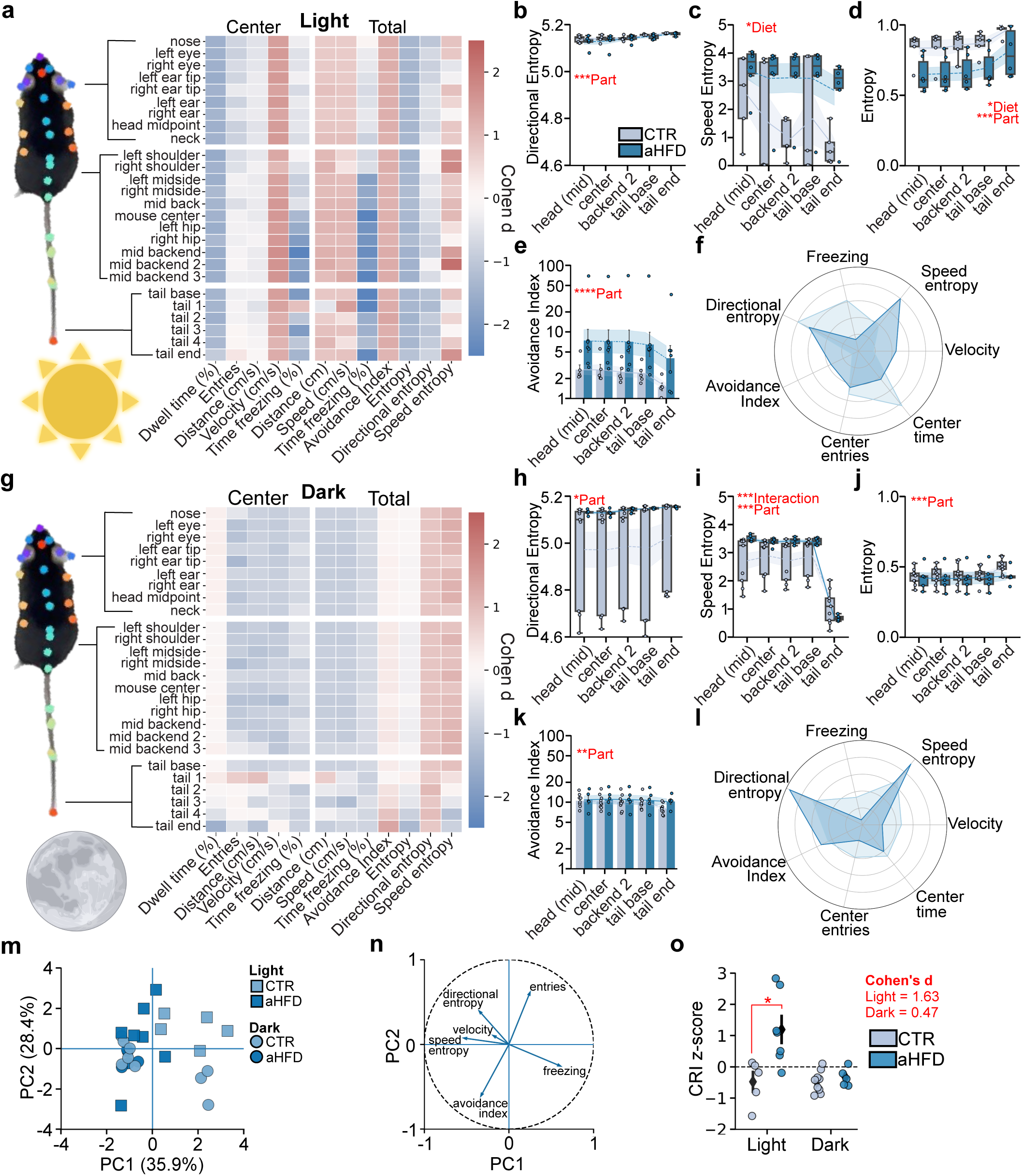
aHFD shifts overall behavioral phenotype in male mice. **(a)** Heatmap of body part specific behavioral changes relative to CTR mice for OFT metrics during the light period in aHFD male mice. **(b)** Mean directional entropy, **(c)** speed entropy, **(d)** overall entropy, and **(e)** avoidance index separated by select body parts. **(f)** Radar plot displaying overall behavioral shift of OFT metrics in CTR and aHFD mice during the light period. **(g)** Heatmap of body part specific behavioral changes relative to CTR mice for OFT metrics during the dark period in aHFD male mice. **(h)** Mean directional entropy, **(i)** speed entropy, **(j)** entropy, and **(k)** avoidance index separated by select body parts. **(l)** Radar plot displaying overall behavioral shift of OFT metrics in CTR and aHFD mice during the dark period. **(m)** PCA analysis showing PC1 against PC2, and **(n)** Top loadings contributing to PCA variance (**Table S5**). **(o)** PCA-loading weighted chronometabolic risk index (CRI) separated by time-of-day (**Table S2**). *Mixed model ANOVA for main effects of diet, body part, and interaction (**Table S3**). CRI scores were analyzed by diet x time two-way ANOVA*. *\*p < 0.05, **p < 0.01, ***p < 0.001*.

### aHFD shifts overall behavioral phenotype in female mice

As described for male mice, we analyzed whole-body movement to assess the broader pattern of behavioral variation in females across the light and dark period (**Fig. 3a,g**). aHFD did not impact directional entropy or overall entropy, but shifted speed entropy, which was primarily attributed to increased posterior-body movement, suggesting more variation in speed localized to the hindlimb region (**Fig. 3b-e; Table S4**). In line with our findings from **Figure 1**, female mice did not demonstrate a change in avoidance index during the light period (**Fig. 3e**), and behavioral shifts were primarily driven by speed entropy and freezing behavior (**Fig. 3f**). aHFD also did not impact entropy measures during the dark period (**Fig. 3h-j**), but interestingly, when examining individual body parts, there was a robust decrease in speed entropy at the end of the tail, indicating more static movement than the rest of the body (**Fig. 3i**). For the avoidance index, values varied by body part, but neither the diet main effect nor the diet × body part interaction reached significance during the dark period (**Fig. 3k; Table S4**). Our behavioral radar plot for the dark period complements our findings in that the mild shifts in time spent in center, entry counts, and freezing time (**Fig. 1f,g**) are the primary drivers of behavioral change and contribute to the overall outcome in avoidance index (**Fig. 3l**). For a behavioral composite, PCA analysis revealed that PC1 (45.1%) and PC2 (29.1%) accounted for 74.2% of variance between CTR and aHFD mice (**Fig. 3m**). Across the top six loadings, PC1 was driven primarily by avoidance index, entry counts, and directional entropy, while PC2 was driven by velocity, freezing time, and entry counts (**Fig. 3n; Table S5**). The exploratory CRI revealed a main diet effect, but did not differ between CTR and aHFD females at either time point after post hoc correction for simple effects (**Fig. 3o**).

**Figure 3.**
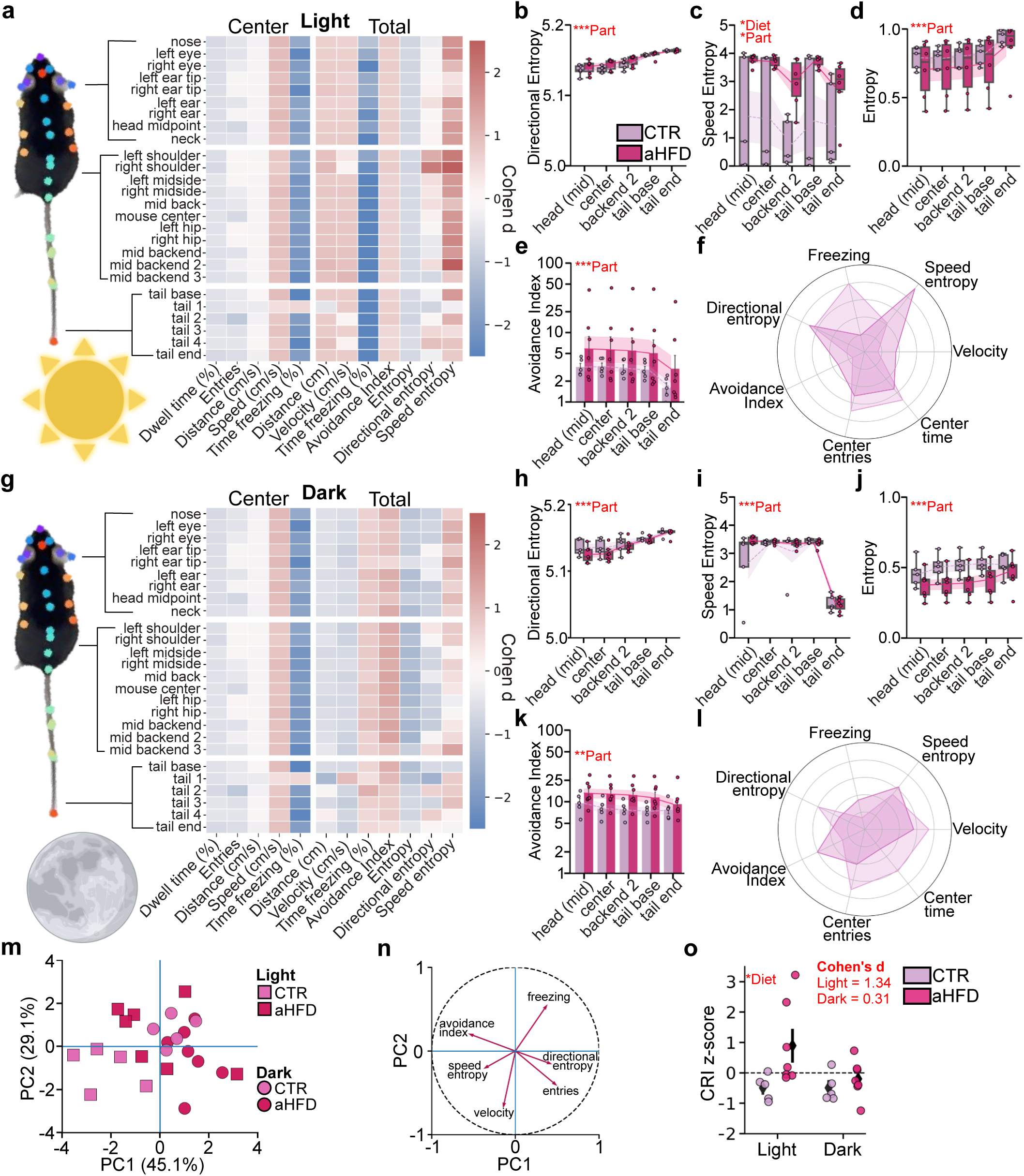
aHFD shifts overall behavioral phenotype in female mice. **(a)** Heatmap of body part specific behavioral changes relative to CTR mice for OFT metrics during the light period in aHFD female mice. **(b)** Mean directional entropy, **(c)** speed entropy, (**d**) overall entropy, and **(e)** avoidance index separated by select body parts. **(f)** Radar plot displaying overall behavioral shift of OFT metrics in CTR and aHFD mice during the light period. **(g)** Heatmap of body part specific behavioral changes relative to CTR mice for OFT metrics during the dark period in aHFD female mice. **(h)** Mean directional entropy, **(i)** speed entropy, **(j)** entropy, and **(k)** avoidance index separated by select body parts. **(l)** Radar plot displaying overall behavioral shift of OFT metrics in CTR and aHFD mice during the dark period. **(m)** PCA analysis showing PC1 against PC2, and **(n)** Top loadings contributing to PCA variance (**Table S5**). **(o)** PCA-loading weighted chronometabolic risk index (CRI) separated by time-of-day (**Table S2**). *Mixed-model ANOVA for main effects of diet, body part, and interaction **(Table S4)***. *CRI scores were analyzed by diet x time two-way ANOVA*. *\*p < 0.05, **p < 0.01, ***p < 0.001*.

### aHFD alters basal properties of layer 2/3 plPFC pyramidal neurons in male and female mice

Circadian and metabolic disruptions are tightly intertwined and influence affective behavior and cognition. Layer 2/3 pyramidal neurons in the plPFC are rhythmic, contribute to cognitive and affective function, and help drive feeding behavior. To unravel how aHFD impacts the physiological function of plPFC layer 2/3 pyramidal neurons, we employed brain slice patch-clamp electrophysiology to test how aHFD influences time-of-day differences in the basal properties of these neurons in male and female mice (**Fig. 4a**). We measured resting membrane potential (RMP), membrane resistance (Rm), and membrane capacitance (Cm) at two different ZT bins (ZT): 6-10, and 18-22. In male mice, aHFD resulted in a depolarization of these neurons, which was limited to the dark period (ZT18-22; **Fig. 4b,c**). This was accompanied by a decrease in membrane capacitance and an increase in membrane resistance (**Fig. 4d,e; Tables S6,S7**), consistent with phase-specific changes in resting membrane conductance in aHFD male mice. In female mice, there was an interaction between diet and ZT bin, but it was not robust enough to drive simple effects (**Fig. 4b,f; Tables S6,S7**) and this was complemented by no change in membrane capacitance, and no change in membrane capacitance or membrane resistance (**Fig. 4g,h**).

**Figure 4.**
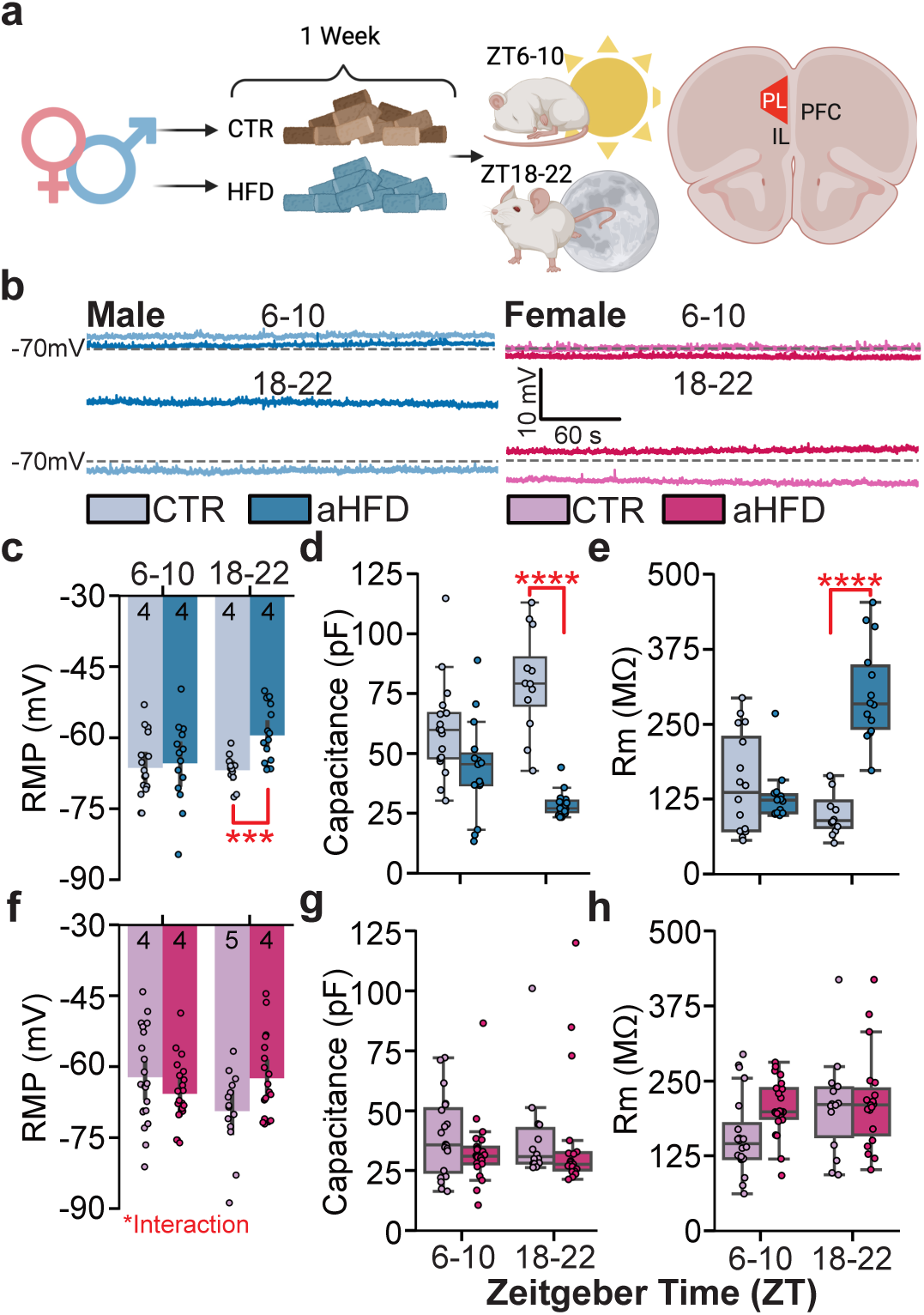
aHFD alters basal properties of plPFC neurons in a time-of-day dependent manner. **(a)** Diagram of experimental approach, ZT bins, and recording location of plPFC layer 2/3 pyramidal neurons**. (b)** Representative traces of RMP in male (*left*) and female (*right*) mice at ZT 6-10 (*top*) and 18-22 (*bottom*). **(c)** Barplot of resting membrane potential (RMP), **(d)** boxplot of membrane capacitance (Cm), and **(e)** membrane resistance (Rm) in male mice. **(f)** Barplot of RMP, and **(g)** boxplot of Cm, and **(h)** Rm in female mice. Numbers inset in ***c,f*** represent animal N values. *Two-way ANOVA for main effect of diet, ZT bin, and interaction **(Tables S6, S7)**. *p < 0.05, **p < 0.01, ***p < 0.001, ****p < 0.0001*.

### aHFD alters action potential dynamics of layer 2/3 plPFC pyramidal neurons in a time-of-day dependent manner in male mice

To determine how changes in intrinsic membrane properties may translate to neuronal output, we next measured evoked action potential firing rate in response to increasing current injections at ZT6-10 and ZT18-22 (**Fig. 5a,b**). In both ZT bins, aHFD did not alter the percentage of layer 2/3 plPFC pyramidal neurons that fired in response to current injections as determined by a Fisher’s Exact test (**Fig. 5c,d**). At ZT6–10, firing curves largely overlapped between diet groups (**Fig. 5c**). At ZT18–22, neurons from aHFD male mice displayed a larger initial increase in firing frequency, reaching a peak 60–80 pA above rheobase, followed by progressive failure to fire at subsequent current steps (**Fig. 5d**). Action potential threshold was unchanged; however, rheobase showed a diet × ZT interaction and was lower in aHFD neurons at ZT18–22, indicating a phase-specific reduction in current required to evoke the first action potential (**Fig. 5e,f**). To determine if these changes in firing properties were accompanied by changes in action potential characteristics, we measured action potential specific parameters such as, half-width, decay tau, afterhyperpolarization (AHP), and total area under the curve (**Fig. 5g-k**). There was no effect of diet or ZT bin on the majority of action potential parameters, apart from aHFD decreasing the total area under the curve (**Fig. 5k; Tables S6,S7**). Seemingly non-significant effects can sometimes compound into a notable shift in physiology. We performed PCA analysis to determine whether the combined changes in basal properties and action potential characteristics reflected broader alterations in the physiological state of layer 2/3 plPFC pyramidal neurons (**Fig. 5l,m**). PC1 and PC2 accounted for 52.1% of the total variance (PC1 = 30.6%, PC2 = 21.5%), with primary loadings presented in **Table S8**. Neurons recorded during the dark period (ZT18-22) showed higher separation between CTR and aHFD than those recorded during the light period (ZT6-10), which were more clustered towards the center, suggesting aHFD produces a more pronounced effect during the dark period (**Fig. 5m**).

**Figure 5.**
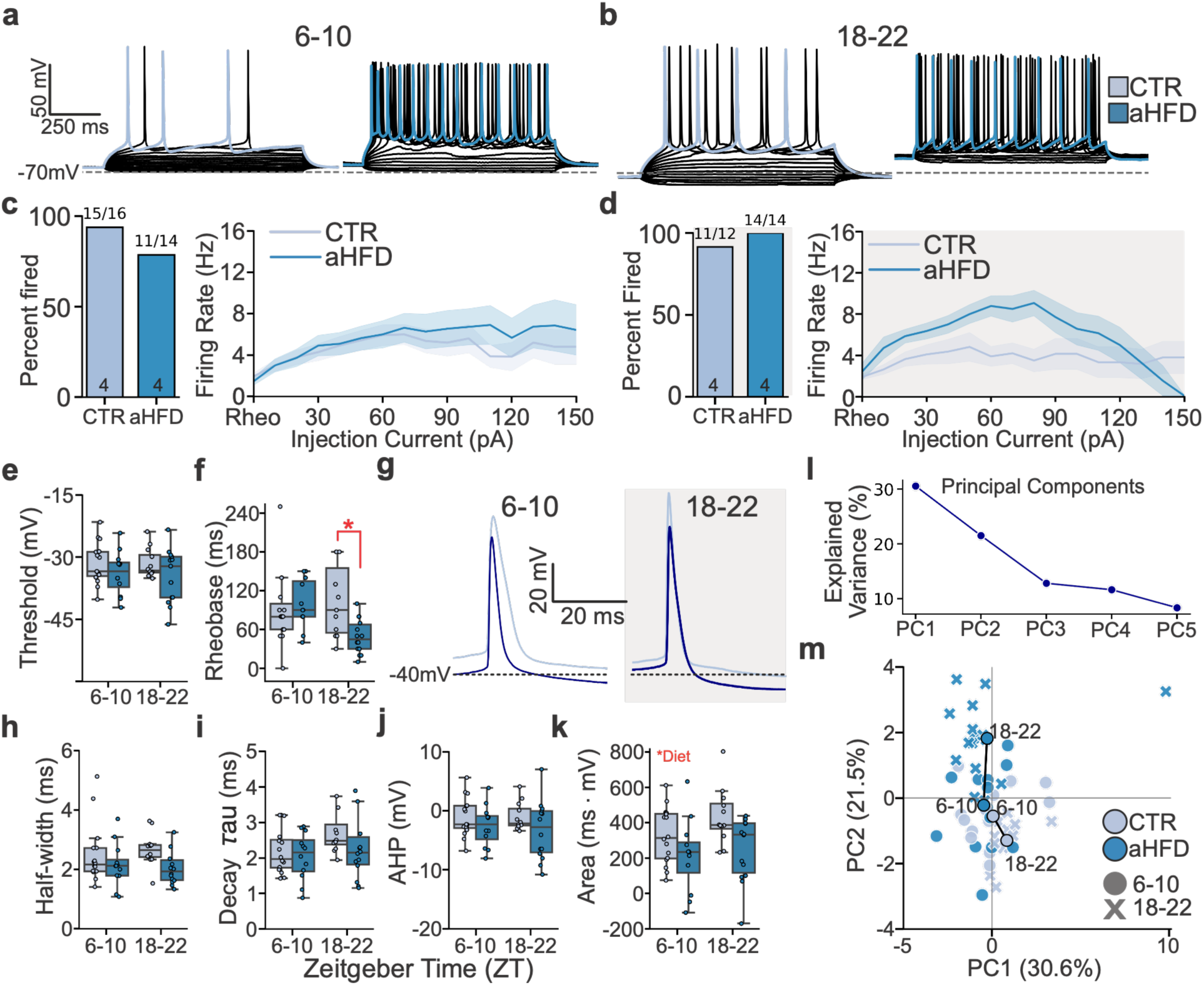
aHFD alters action potential dynamics in plPFC L2/3 pyramidal neurons of male mice. **(a)** Representative traces of evoked action potentials in CTR and aHFD male mice at ZT6-10, and **(b)** ZT18-22. **(c)** Percent of neurons that fired with current clamp injections (*left*) and firing rates with 10 pA current steps from rheobase at ZT6-10 and **(d)** 18-22. **(e)** Boxplot of action potential threshold, and **(f)** rheobase. **(g)** Representative trace of individual action potentials, and **(h)** boxplot of half-width, **(i)** decay tau, **(j)** afterhyperpolarization (AHP), and **(k)** total action potential area. **(l)** PCA analysis inclusive of action potential characteristics in CTR and aHFD with explained variance for top 5 principal components (PC), and **(m)** point plot of variance attributed to PC1 and PC2, with centroids showing mean values separated by ZT bin. Numbers inset in ***c,d*** represent animal N values. *Two-way ANOVA for the main effect of diet, ZT bin, and interaction **(Tables S6-8)**. Sidak post hoc for simple effects. *p < 0.05*.

### aHFD alters action potential dynamics of layer 2/3 plPFC pyramidal neurons in female mice

We performed the same analyses in female mice to determine whether aHFD altered action potential dynamics. Evoked firing curves showed substantial overlap between diet groups, and the proportion of responsive neurons did not differ at either ZT bin (Fisher’s exact test: ZT6–10, *p* = 1.00; ZT18–22, *p* = 0.106; **Fig. 6a–d**). Thus, effects of aHFD on female firing rates were minimal.

**Figure 6.**
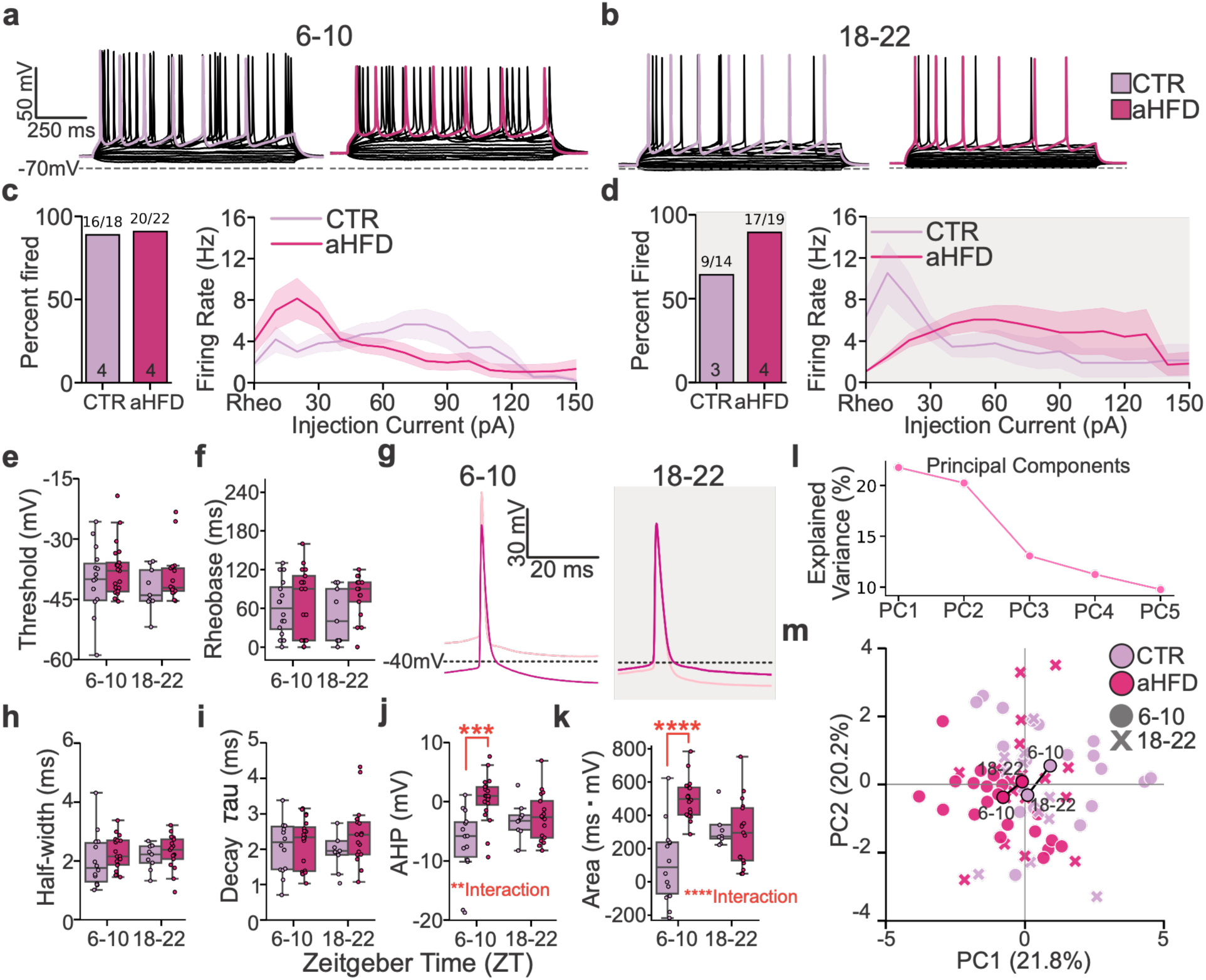
aHFD alters action potential dynamics in plPFC L2/3 pyramidal neurons in female mice. **(a)** Representative traces of evoked action potentials in CTR and aHFD female mice at ZT6-10, and **(b)** ZT18-22. **(c)** Percent of neurons that fired with current clamp injections (*left*) and firing rates with 10pA current steps from rheobase at ZT6-10 and **(d)** 18-22. **(e),** Boxplot of action potential threshold, and **(f)** rheobase. **(g)** Representative trace of individual action potentials, and **(h)** boxplot of half-width, **(i)** decay tau, **(j)** afterhyperpolarization (AHP), and **(k)** total action potential area. **(l)** PCA analysis inclusive of action potential characteristics in CTR and aHFD with explained variance for top 5 PCs, and **(m)** point plot showing individual and mean values separated by ZT bin. Number inserts in ***c,d*** represent animal N values. *Two-way ANOVA for the main effect of diet, ZT, and interaction **(Tables S6-8)**. Sidak post hoc for simple effects. **p < 0.01, ***p < 0.001, ****p < 0.0001*.

We next examined whether these firing profiles were accompanied by alterations in action potential characteristics. Action potential threshold, rheobase, half-width, and decay tau did not differ by diet or ZT (**Fig. 6e–i; Tables S6,S7**). In contrast, AHP amplitude and action potential area showed diet and diet × ZT effects, with attenuated AHPs and increased action potential area in aHFD neurons appearing most pronounced at ZT6-10 (**Fig. 6j,k**). Thus, aHFD did not broadly alter action potential threshold or kinetics in females but selectively affected waveform dynamics in a time-of-day dependent manner.

To determine whether combined intrinsic electrophysiological properties distinguished neurons by treatment and time-of-day, we performed PCA using the measured basal and action potential characteristics (**Fig. 6l,m**). PC1 (21.8%) and PC2 (20.2%) explained 42.0% of total variance. Although substantial overlap remained, neurons from aHFD mice showed greater dispersion across PC space at ZT6–10 (**Fig. 6m**), suggesting a stronger multivariate distinction during the light period.

### Sex-specific effects of aHFD on the current-voltage relationship are time-of-day dependent

To determine whether aHFD altered voltage-dependent membrane currents, we normalized steady-state current to membrane capacitance and compared derived current–voltage measures at ZT6–10 and ZT18–22 (**Fig. 7**). In males, depolarized-range slope conductance showed a diet × ZT interaction, with greater conductance in aHFD neurons at ZT18–22 (**Fig. 7g,h**). Neither hyperpolarized-range slope conductance nor apparent reversal potential differed by diet or ZT (**Fig. 7b,e,f**). In females, hyperpolarized-range slope conductance did not differ by diet or ZT, and there was no interaction (**Fig. 7i,j**). Apparent reversal potential also showed effects of ZT and a diet × ZT interaction and shifted negatively in aHFD neurons from female mice at ZT6–10 (**Fig. 7c,d**). Depolarized-range slope conductance did not differ in females, although the interaction approached significance (**Fig. 7k,l**). Together, these results indicate sex specific and time-of-day dependent changes in net membrane conductance following aHFD.

**Figure 7.**
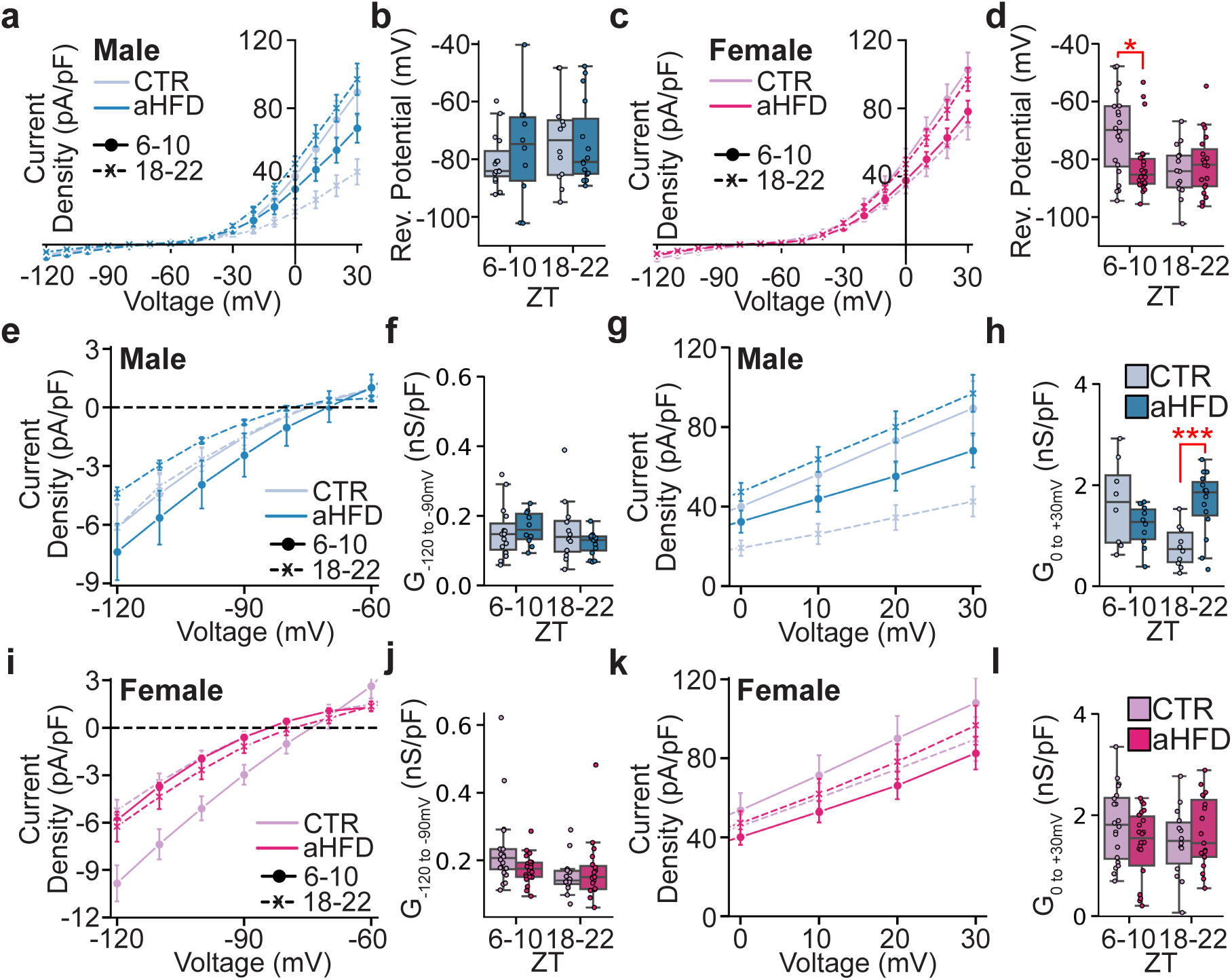
aHFD alters conductance of layer 2/3 pyramidal neurons in a time-of-day dependent manner. **(a)** I-V curve line plot of male and **(c)** female current density in CTR and aHFD mice at ZT6-10 and 18-22. **(b)** Apparent reversal potential calculated from I-V curves in male and **(d)** female mice. **(e)** Current normalized to capacitance (current density) plot of hyperpolarized conductance (*G*), and **(f)** calculated conductance density from -120mV to -90mV in male mice. **(g)** Current density plot of depolarized conductance, and **(h)** calculated conductance from 0mV to +30mV in male mice. **(i)** Current density plot of hyperpolarized conductance (G), and **(j)** calculated conductance density from -120mV to -90mV in female mice. **(k)** Current density plot of depolarized conductance, and **(l)** calculated conductance density from 0mV to +30mV in female mice. *Two-way ANOVA for the main effect of diet, ZT bin, and interaction **(Tables S6, S7)**. Sidak post hoc for simple effects. *p < 0.05, **p < 0.01*.

## DISCUSSION

Obesity and metabolic disease have become a long-standing epidemic in the United States^2,52^. An HFD not only has long-term detrimental effects on health but can perturb physiological function and behavior before the onset of obesity. Independent of obesity, an HFD can also produce rapid consequences for brain function, with cognitive and mood-related deficits emerging long before signs of obesity or related systemic complications appear. While prior work has shown that HFD induces anxiety-like behavior and cognitive changes in mice^5^, many unanswered questions remain in our understanding of how short-term HFD impacts affective behaviors and PFC function. Here we address this gap with three main findings. First, one-week aHFD exposure altered behavior and locomotion, which was sex specific and time-of-day dependent. Second, aHFD altered intrinsic properties and firing profiles in a phase-specific manner. Third, current–voltage analysis identified sex specific time-of-day dependent shifts in apparent reversal potential and membrane conductance.

Previous work has demonstrated that behavioral performance is influenced by circadian phase and has emphasized the importance of considering time-of-day and light/dark cycles when interpreting behavioral assays^9,53^. Several studies have also shown that short-term HFD exposure is sufficient to induce anxiety-like behavior, though the earliest reported effects emerged 2-3 weeks after exposure^5,49^. Building on this framework, our behavioral studies imply that aHFD produces an anxiety-like state expressed through active avoidance and increased but less structured movement in males evident after only one week of HFD exposure. These effects primarily occurred in the inactive period, a period where controls were most exploratory and least avoidant. The diet effect emerging in this phase carries several implications. It indicates that acute HFD does not simply raise anxiety-like behavior overall but narrows the diurnal window in which exploratory behavior is normally least constrained, flattening the behavioral contrast between the active and inactive periods. Females showed a comparable diurnal organization of baseline behavior but a weaker response to diet, with reduced freezing during the inactive phase and no significant diet effect on the avoidance index. Two implications follow. First, the behavioral consequences of aHFD and the impact of time-of-day are sex-specific. Second, the absence of a strong effect on these measures does not establish that female plPFC function is spared; instead, it indicates that female responses to acute dietary perturbation may unfold differently from males. The exploratory CRI showed its clearest male separation during the light period, whereas neither female post hoc comparison reached significance. Together, these findings extend previous work by demonstrating that aHFD disrupts the normal diurnal organization of exploratory and affective behaviors and these effects may be sex specific. However, a formal and properly controlled sex differences study would be needed to determine the differences in male and female behavior.

In addition to demonstrating how short-term exposure to HFD affects behavior, we show that it also disrupts normal diurnal differences in neuronal excitability. While previous work^23,24^ found that plPFC pyramidal neurons are normally hyperpolarized during the active phase (dark period), requiring stronger input current to fire, we observed the opposite after aHFD: a depolarization during the dark period. In males, this depolarization was accompanied by an increased membrane resistance and reduced capacitance during the active phase, consistent with altered resting membrane conductance rather than identifying a specific channel. This interpretation is supported by the precedent that many neuronal populations in the plPFC are diurnally modulated, with disruptions in circadian rhythms being detrimental for cognitive and emotional health^23,25^. The current–voltage analysis further showed increased depolarized-range slope conductance in males at ZT18–22, arguing against a simple global loss of potassium (K^+^) conductance. Together, these changes could make neurons easier to recruit at lower inputs while limiting sustained firing during stronger depolarization. One possibility is that the depolarized resting state and altered voltage-dependent conductance increase sodium-channel inactivation during prolonged current injection; direct channel-isolation experiments will be required to test this mechanism. These findings suggest that aHFD alters how plPFC pyramidal neurons receive and transmit signals in a time-of-day-dependent manner.

The current–voltage data provide candidate mechanisms rather than channel identity. The greater depolarized-range conductance density in aHFD males at ZT18–22 is consistent with recruitment of one or more outward currents that could constrain sustained firing. Kv1.3 is one metabolism-associated candidate^54,55^, but the present mixed whole-cell currents cannot distinguish Kv1.3 from other voltage- or calcium-activated K⁺ conductances. Additionally, HCN, Kir2, or K⁺ leak channels warrant investigation as potential contributors to changes in PFC pyramidal neuron excitability given that they jointly shape frontal pyramidal-neuron excitability^56–58^ and influence rhythms in membrane physiology^59^. Pharmacological isolation and current-kinetic analyses will be required to test these possibilities.

Beyond changes in intrinsic membrane properties, an important next step will be to determine how aHFD affects excitatory/inhibitory (E/I) balance within the plPFC. Our data demonstrate sex-specific and time-of-day dependent changes in intrinsic excitability and membrane conductance, but whether this reflects enhanced glutamatergic drive, reduced GABAergic inhibition, or both, remains unresolved. This question is also temporally relevant because synaptic inhibition varies across the 24-hour cycle in pyramidal neurons of the prefrontal cortex and other forebrain regions^25,60^. Prior work in the lateral orbitofrontal cortex (lOFC) demonstrated that even seven days of HFD increases pyramidal neuron excitability in association with reduced tonic GABAergic inhibition, whereas prolonged diet exposure produces additional deficits in synaptic inhibitory transmission^28^. Another study of diet-induced obesity similarly reported depolarized resting membrane potential and reduced inhibitory synaptic transmission onto lOFC pyramidal neurons, attributable to decreased GABA release probability^61^. Whether comparable disinhibition occurs in plPFC, and if it is accompanied by changes in glutamatergic drive, remains untested. While defining E/I balance will clarify the synaptic mechanisms underlying aHFD-induced excitability, an equally important question is how these physiological changes alter communication within larger metabolic-cognitive circuits.

In summary, here we sought to determine the extent to which acute dietary perturbation alters cortical function in the period before obesity emerges, specifically after just one week of exposure. We achieved this through behavioral and physiological approaches, showing that diet alone can alter plPFC intrinsic properties, firing, and membrane conductance in a time-of-day dependent manner, alongside changes in behavior. In doing so, we address a critical gap in the literature, highlighting how diet and time-of-day are closely intertwined in mediating plPFC function, and associated behavioral outputs. Through these approaches, we address the underexplored question of how aHFD consumption impacts the brain, demonstrating that behavior and prefrontal physiology are vulnerable to disruption within a remarkably short window of exposure. Together, these findings indicate that the PFC is an early neural target of dietary perturbation, with aHFD disrupting physiology and behavior across multiple levels of organization.

## ACKNOWLEDGEMENTS

This work was supported by National Institutes of Health (NIH) NIGMS COBRE 5P20GM121310, NIH NIGMS INBRE 2P20GM103432, and startup funding to BLR from the University of Wyoming. We thank Robert M. Carroll for his exceptional vivarium management and invaluable support and expertise in animal care and husbandry throughout this study.

## CONTRIBUTIONS

BLR conceptualized the project. BLR, PPW, and NCC contributed to data collection, analysis, and manuscript writing. JKC and JET contributed to data collection. WJW contributed to data collection, editing of manuscript, and project oversight.

## DATA AND CODE AVAILABILITY

The data for this study is publicly available on Zenodo with DOI: https://doi.org/10.5281/zenodo.22859974. The Zenodo package also includes all analysis code and Python notebooks for reproducing all quantitiative figures and statistical analysis. Raw acquisition-level data and are available upon request.

## Declaration of generative AI and AI-assisted technologies in the manuscript preparation process

During the preparation of this work, a subset of the authors used OpenAI ChatGPT 5.6 and 6.0 to facilitate locating primary literature, cross-checking consistency of statistics among figures, text, tables, methods, and the original output code. AI was also used to troubleshoot, clean, and organize analysis code. All drafts, analysis, concepts, edits, figures, original code, and statistics were author generated. All AI facilitated tasks and edits were manually vetted and incorperated by the authors. Final edits and preparation for submission was performed by the authors. All AI use was performed locally and no components of this manuscript were used for model training. After using this tool, the authors reviewed and edited the content as needed and take full responsibility for the content of the published article.

## REFERENCES

1. CDC. Obesity is a Common, Serious, and Costly Disease. Centers for Disease Control and Prevention https://www.cdc.gov/obesity/data/adult.html (2022).

2. Ng, M. et al. National-level and state-level prevalence of overweight and obesity among children, adolescents, and adults in the USA, 1990–2021, and forecasts up to 2050. The Lancet 404, 2278–2298 (2024).

3. Wali, J. A. et al. Cardio-Metabolic Effects of High-Fat Diets and Their Underlying Mechanisms—A Narrative Review. Nutrients 12, 1505 (2020).

4. Evans, A. K. et al. Impact of high-fat diet on cognitive behavior and central and systemic inflammation with aging and sex differences in mice. Brain. Behav. Immun. 118, 334–354 (2024).

5. Gainey, S. J. et al. Short-Term High-Fat Diet (HFD) Induced Anxiety-Like Behaviors and Cognitive Impairment Are Improved with Treatment by Glyburide. Front. Behav. Neurosci. 10, (2016).

6. Baker, K. D. & Reichelt, A. C. Impaired fear extinction retention and increased anxiety-like behaviours induced by limited daily access to a high-fat/high-sugar diet in male rats: Implications for diet-induced prefrontal cortex dysregulation. Neurobiol. Learn. Mem. 136, 127–138 (2016).

7. de Paula, G. C. et al. Hippocampal Function Is Impaired by a Short-Term High-Fat Diet in Mice: Increased Blood-Brain Barrier Permeability and Neuroinflammation as Triggering Events. Front. Neurosci. 15, 734158 (2021).

8. Daut, R. A. et al. Circadian misalignment has differential effects on affective behavior following exposure to controllable or uncontrollable stress. Behav. Brain Res. 359, 440–445 (2019).

9. Karatsoreos, I. N., Bhagat, S., Bloss, E. B., Morrison, J. H. & McEwen, B. S. Disruption of circadian clocks has ramifications for metabolism, brain, and behavior. Proc. Natl. Acad. Sci. 108, 1657–1662 (2011).

10. Karatsoreos, I. N. Effects of circadian disruption on mental and physical health. Curr Neurol Neurosci Rep 12, 218–25 (2012).

11. van der Vinne, V., Martin Burgos, B., Harrington, M. E. & Weaver, D. R. Deconstructing circadian disruption: Assessing the contribution of reduced peripheral oscillator amplitude on obesity and glucose intolerance in mice. J. Pineal Res. e12654 (2020) doi:10.1111/jpi.12654.

12. Reppert, S. M. & Weaver, D. R. Coordination of circadian timing in mammals. Nature 418, 935–941 (2002).

13. Landgraf, D. et al. Genetic Disruption of Circadian Rhythms in the Suprachiasmatic Nucleus Causes Helplessness, Behavioral Despair, and Anxiety-like Behavior in Mice. Biol. Psychiatry 80, 827–835 (2016).

14. Roybal, K. et al. Mania-like behavior induced by disruption of CLOCK. Proc. Natl. Acad. Sci. U. S. A. 104, 6406–6411 (2007).

15. Logan, R. W. & McClung, C. A. Rhythms of life: circadian disruption and brain disorders across the lifespan. Nat. Rev. Neurosci. 20, 49–65 (2019).

16. Williams, D. L. & Schwartz, M. W. Out of synch: *Clock* mutation causes obesity in mice. Cell Metab. 1, 355–356 (2005).

17. Miller, E. K. & Cohen, J. D. An Integrative Theory of Prefrontal Cortex Function. Annu. Rev. Neurosci. 24, 167–202 (2001).

18. McEwen, B. S. & Morrison, J. H. Brain On Stress: Vulnerability and Plasticity of the Prefrontal Cortex Over the Life Course. Neuron 79, 16–29 (2013).

19. Yuen, E. Y. et al. Acute stress enhances glutamatergic transmission in prefrontal cortex and facilitates working memory. Proc. Natl. Acad. Sci. 106, 14075–14079 (2009).

20. Hains, A. B. & Arnsten, A. F. T. Molecular mechanisms of stress-induced prefrontal cortical impairment: implications for mental illness. Learn. Mem. Cold Spring Harb. N 15, 551–564 (2008).

21. Fu, C.-C. et al. PPARγ Dysfunction in the Medial Prefrontal Cortex Mediates High-Fat Diet-Induced Depression. Mol. Neurobiol. 59, 4030–4043 (2022).

22. Saitoh, A. et al. Activation of the prelimbic medial prefrontal cortex induces anxiety-like behaviors via N-Methyl-D-aspartate receptor-mediated glutamatergic neurotransmission in mice. J. Neurosci. Res. 92, 1044–1053 (2014).

23. Roberts, B. L. & Karatsoreos, I. N. Circadian desynchronization disrupts physiological rhythms of prefrontal cortex pyramidal neurons in mice. Sci. Rep. 13, 9181 (2023).

24. Roberts, B. L. et al. Perinatal circadian desynchronization disrupts sleep and prefrontal cortex function in adult offspring. Sleep 49, zsaf210 (2026).

25. Burns, J. N. et al. Molecular and cellular rhythms in excitatory and inhibitory neurons in the mouse prefrontal cortex. Cereb. Cortex 35, bhaf188 (2025).

26. Martínez-Orozco, H., Reyes-Castro, L. A., Lomas-Soria, C., Solís-Ortíz, S. & Diaz-Miranda, S. Y. Prefrontal Cortex Dysregulation of Amino Acid-Glucose Homeostasis Links High-Fat and/or High-Fructose Intake to Cognitive Deficits in Male Mice. Neurochem. Res. 51, 184 (2026).

27. Huang, H. et al. A Four-Week High-Fat Diet Induces Anxiolytic-like Behaviors through Mature BDNF in the mPFC of Mice. Brain Sci. 14, 389 (2024).

28. Seabrook, L. T. et al. Short- and Long-Term High-Fat Diet Exposure Differentially Alters Phasic and Tonic GABAergic Signaling onto Lateral Orbitofrontal Pyramidal Neurons. J. Neurosci. Off. J. Soc. Neurosci. 43, 8582–8595 (2023).

29. Atar, A. Neurobiological Consequences of High-Fat High-Sugar Diets on the Mesocorticolimbic System: a Narrative Review. Curr. Nutr. Rep. 15, 6 (2026).

30. Khazen, T., Hatoum, O. A., Ferreira, G. & Maroun, M. Acute exposure to a high-fat diet in juvenile male rats disrupts hippocampal-dependent memory and plasticity through glucocorticoids. Sci. Rep. 9, 12270 (2019).

31. Kanoski, S. E., Zhang, Y., Zheng, W. & Davidson, T. L. The effects of a high-energy diet on hippocampal function and blood-brain barrier integrity in the rat. J. Alzheimers Dis. JAD 21, 207–219 (2010).

32. Walf, A. A. & Frye, C. A. The use of the elevated plus maze as an assay of anxiety-related behavior in rodents. Nat. Protoc. 2, 322–328 (2007).

33. Seibenhener, M. L. & Wooten, M. C. Use of the Open Field Maze to Measure Locomotor and Anxiety-like Behavior in Mice. J. Vis. Exp. JoVE 52434 (2015) doi:10.3791/52434.

34. Mathis, A. et al. DeepLabCut: markerless pose estimation of user-defined body parts with deep learning. Nat. Neurosci. 21, 1281–1289 (2018).

35. Tanaka, S., Young, J. W., Halberstadt, A. L., Masten, V. L. & Geyer, M. A. Four factors underlying mouse behavior in an open field. Behav. Brain Res. 233, 10.1016/j.bbr.2012.04.045 (2012).

36. Prut, L. & Belzung, C. The open field as a paradigm to measure the effects of drugs on anxiety-like behaviors: a review. Eur. J. Pharmacol. 463, 3–33 (2003).

37. Ye, S. et al. SuperAnimal pretrained pose estimation models for behavioral analysis. Nat. Commun. 15, 5165 (2024).

38. Shannon, C. E. A Mathematical Theory of Communication. Bell Syst. Tech. J. 27, 379–423 (1948).

39. Moore, T. Y., Cooper, K. L., Biewener, A. A. & Vasudevan, R. Unpredictability of escape trajectory explains predator evasion ability and microhabitat preference of desert rodents. Nat. Commun. 8, 440 (2017).

40. Paulus, M. P., Geyer, M. A., Gold, L. H. & Mandell, A. J. Application of entropy measures derived from the ergodic theory of dynamical systems to rat locomotor behavior. Proc. Natl. Acad. Sci. U. S. A. 87, 723–727 (1990).

41. Ting, J. T. et al. Preparation of Acute Brain Slices Using an Optimized N-Methyl-D-glucamine Protective Recovery Method. JoVE J. Vis. Exp. e53825 (2018) doi:10.3791/53825.

42. Pedregosa, F. et al. Scikit-learn: Machine Learning in Python. J. Mach. Learn. Res. 12, 2825–2830 (2011).

43. Jolliffe, I. T. & Cadima, J. Principal component analysis: a review and recent developments. Philos. Transact. A Math. Phys. Eng. Sci. 374, 20150202 (2016).

44. Guilloux, J.-P., Seney, M., Edgar, N. & Sibille, E. Integrated behavioral z-scoring increases the sensitivity and reliability of behavioral phenotyping in mice: relevance to emotionality and sex. J. Neurosci. Methods 197, 21–31 (2011).

45. Cohen, J. Statistical Power Analysis for the Behavioral Sciences. (Routledge, New York, 2013). doi:10.4324/9780203771587.

46. Roberts, B. L., Bennett, C. M., Carroll, J. M., Lindsley, S. R. & Kievit, P. Early overnutrition alters synaptic signaling and induces leptin resistance in arcuate proopiomelanocortin neurons. Physiol. Behav. 206, 166– 174 (2019).

47. Hasler, G. et al. The associations between psychopathology and being overweight: a 20-year prospective study. Psychol. Med. 34, 1047–1057 (2004).

48. Mohanty, P. et al. Metabolic Dysfunction in Psychiatric Disorders: A Narrative Review of Shared Pathways Between Mental Illness and Cardiometabolic Disease. Cureus 18, e111469.

49. Machado, A. E., Rezer, P., Mancini, G., Latini, A. & Moreira, E. L. G. Short-term high-fat diet alters behavior, peripheral metabolism, and brain mitochondrial function in Swiss mice. An. Acad. Bras. Ciênc. 96, e20240880 (2024).

50. Shansky, R. M. Behavioral neuroscience’s inevitable SABV growing pains. Trends Neurosci. 47, 669–676 (2024).

51. Walton, J. C., Bumgarner, J. R. & Nelson, R. J. Sex Differences in Circadian Rhythms. Cold Spring Harb. Perspect. Biol. 14, a039107 (2022).

52. Kranjac, A. W. & Kranjac, D. Explaining adult obesity, severe obesity, and BMI: Five decades of change. Heliyon 9, e16210 (2023).

53. Tsao, C.-H., Flint, J. & Huang, G.-J. Influence of diurnal phase on behavioral tests of sensorimotor performance, anxiety, learning and memory in mice. Sci. Rep. 12, 432 (2022).

54. Al Koborssy, D. et al. Modulation of olfactory-driven behavior by metabolic signals: role of the piriform cortex. Brain Struct. Funct. 224, 315–336 (2019).

55. Xu, J. et al. The voltage-gated potassium channel Kv1.3 regulates energy homeostasis and body weight. Hum. Mol. Genet. 12, 551–559 (2003).

56. Day, M. et al. Dendritic excitability of mouse frontal cortex pyramidal neurons is shaped by the interaction among HCN, Kir2, and Kleak channels. J. Neurosci. Off. J. Soc. Neurosci. 25, 8776–8787 (2005).

57. Song, C. & Moyer, J. R. Layer- and subregion-specific differences in the neurophysiological properties of rat medial prefrontal cortex pyramidal neurons. J. Neurophysiol. 119, 177–191 (2018).

58. Bekkers, J. M. Properties of voltage-gated potassium currents in nucleated patches from large layer 5 cortical pyramidal neurons of the rat. J. Physiol. 525 Pt 3, 593–609 (2000).

59. Paul, J. R. et al. Circadian regulation of membrane physiology in neural oscillators throughout the brain. Eur. J. Neurosci. 51, 109–138 (2020).

60. Bridi, M. C. D. et al. Daily Oscillation of the Excitation-Inhibition Balance in Visual Cortical Circuits. Neuron 105, 621–629.e4 (2020).

61. Thompson, J. L. et al. Obesity-Induced Structural and Neuronal Plasticity in the Lateral Orbitofrontal Cortex. Neuropsychopharmacol. Off. Publ. Am. Coll. Neuropsychopharmacol. 42, 1480–1490 (2017).

