## Supplemental Tables for "Brief high-fat diet exposure temporally reshapes affective behavior and prefrontal cortex function"

**Supplemental Table 1. Open-field test mean values for diet and light cycle from mouse center**

| Open Field Test Values Mean (SEM) | | | | |
|---|---|---|---|---|
| **Males** | **Light** | | **Dark** | |
| **Parameter** | **CTR (N = 5)** | **aHFD (N = 7)** | **CTR (N = 9)** | **aHFD (N = 5)** |
| Center Entries | 48.80 (3.31) | 42.86 (9.4) | 32.22 (4.28) | 24.80 (1.36) |
| Center Time (%) | 27.90 (3.38) | 15.1 (3.02) | 9.10 (0.64) | 8.57 (1.26) |
| Center Distance (cm) | 1110.84 (113.88) | 856.67 (195.39) | 618.04 (60.59) | 473.64 (37.27) |
| Speed (cm/s) | 12.08 (0.71) | 15.03 (1.21) | 16.29 (0.98) | 14.07 (0.65) |
| Freezing Time (%) | 3.75 (0.47) | 1.55 (0.23) | 2.73 (0.74) | 1.50 (0.17) |
| Avoidance Index | 2.91 (0.68) | 14.22 (9.00) | 10.47 (0.89) | 11.63 (1.71) |
| Directional Entropy | 5.14 (0.01) | 5.13 (0.01) | 4.97 (0.08) | 5.13 (0.01) |
| Speed Entropy | 1.51 (0.91) | 3.13 (0.44) | 2.88 (0.23) | 3.36 (0.07) |
| Entropy | 0.84 (0.06) | 0.57 (0.09) | 0.44 (0.02) | 0.42 (0.04) |
| CRI z-score | -0.48 (0.31) | 1.18 (0.44) | -0.48 (0.12) | -0.31 (0.14) |
| **Females** | **Light** | | **Dark** | |
| **Parameter** | **CTR (N = 5)** | **aHFD (N = 7)** | **CTR (N = 5)** | **aHFD (N = 7)** |
| Center Entries | 56.20 (7.14) | 45.14 (7.02) | 42.40 (2.94) | 31.29 (5.01) |
| Center Time (%) | 23.86 (2.12) | 18.94 (4.56) | 11.41 (1.39) | 7.59 (1.04) |
| Center Distance (cm) | 1190.40 (142.20) | 1109.28 (215.74) | 820.69 (99.56) | 582.36 (98.14) |
| Speed (cm/s) | 13.44 (0.79) | 15.99 (1.46) | 16.61 (1.01) | 14.94 (1.10) |
| Freezing Time (%) | 3.92 (0.25) | 1.59 (0.23) | 0.99 (0.15) | 1.55 (0.31) |
| Avoidance Index | 3.33 (0.39) | 10.61 (5.71) | 8.30 (1.15) | 13.70 (1.99) |
| Directional Entropy | 5.14 (0.00) | 5.14 (0.01) | 5.14 (0.01) | 5.13 (0.00) |
| Speed Entropy | 1.65 (0.90) | 3.66 (0.09) | 3.38 (0.03) | 3.41 (0.03) |
| Entropy | 0.79 (0.04) | 0.63 (0.11) | 0.51 (0.04) | 0.38 (0.04) |
| CRI z-score | -0.46 (0.18) | 0.93 (0.49) | -0.46 (0.22) | -0.27 (0.25) |

**Supplemental Table 2. Open-field test statistics for diet and light cycle from mouse center**

| Open Field Test Statistics (mouse center) | | | | | | |
|---|---|---|---|---|---|---|
| **Males** | **Diet** | | **Light Cycle** | | **Interaction** | |
| **Parameter** | ***F*** | ***p*** | ***F*** | ***p*** | ***F*** | ***p*** |
| Center Entries | 1.15 | 0.295 | 7.62 | ***0.011*** | 0.01 | 0.907 |
| Center time (%) | 8.13 | 0.009 | 33.66 | ***<0.001*** | 7.54 | ***0.012*** |
| Center distance (cm) | 2.42 | 0.134 | 12.15 | ***0.002*** | 0.19 | 0.669 |
| Velocity (cm/s) | 0.05 | 0.821 | 2.76 | 0.111 | 6.00 | ***0.023*** |
| Freezing time (%) | 8.38 | ***0.008*** | 0.90 | 0.354 | 0.69 | 0.413 |
| Avoidance Index (ln) | 4.32 | ***0.049*** | 13.36 | ***0.001*** | 3.32 | 0.082 |
| Directional entropy | 1.88 | 0.184 | 2.48 | 0.129 | 2.32 | 0.142 |
| Speed entropy | 4.91 | ***0.037*** | 3.25 | 0.085 | 1.52 | 0.231 |
| Entropy | 5.35 | ***0.030*** | 23.80 | ***<0.001*** | 4.25 | 0.051 |
| CRI z-score | 8.78 | ***0.007*** | 5.81 | ***0.025*** | 6.40 | ***0.019*** |
| **Females** | **Diet** | | **Light Cycle** | | **Interaction** | |
| **Parameter** | ***F*** | ***p*** | ***F*** | ***p*** | ***F*** | ***p*** |
| Center entries | 3.32 | 0.083 | 5.32 | ***0.032*** | 0.00 | 0.996 |
| Center time (%) | 2.13 | 0.160 | 15.99 | ***0.001*** | 0.03 | 0.856 |
| Center distance (cm) | 1.00 | 0.328 | 8.63 | ***0.008*** | 0.24 | 0.627 |
| Velocity (cm/s) | 0.13 | 0.722 | 0.35 | 0.559 | 3.01 | 0.098 |
| Freezing time (%) | 11.57 | ***0.003*** | 23.33 | ***<0.001*** | 30.62 | ***<0.001*** |
| Avoidance Index (ln) | 3.52 | 0.075 | 9.71 | ***0.005*** | 0.026 | 0.874 |
| Directional entropy | 0.64 | 0.433 | 3.60 | 0.072 | 1.55 | 0.227 |
| Speed entropy | 7.33 | ***0.014*** | 2.41 | 0.136 | 6.98 | ***0.016*** |
| Entropy | 3.68 | 0.069 | 13.64 | ***0.001*** | 0.03 | 0.858 |
| CRI z-score | 5.06 | ***0.036*** | 4.13 | 0.056 | 2.95 | 0.101 |

**Supplemental Table 3. Male mixed-model ANOVA statistics for diet and body part movement**

| Male Open Field Test Statistics (Diet x Body Part) | | | | | | |
|---|---|---|---|---|---|---|
| **Light Cycle** | **Diet** | | **Body Part** | | **Interaction** | |
| **Parameter** | ***F*** | ***p*** | ***F*** | ***p*** | ***F*** | ***p*** |
| Center entries | 0.16 | 0.696 | 15.63 | ***0.001*** | 0.25 | 0.910 |
| Center time (%) | 7.78 | ***0.019*** | 17.07 | ***0.001*** | 0.35 | 0.844 |
| Center distance (cm) | 0.78 | 0.397 | 24.25 | ***<0.001*** | 0.09 | 0.985 |
| Velocity (cm/s) | 2.53 | 0.143 | 186.72 | ***<0.001*** | 1.30 | 0.288 |
| Freezing time (%) | 15.06 | ***0.003*** | 27.61 | ***0.001*** | 18.48 | ***<0.001*** |
| Avoidance Index (ln) | 4.04 | 0.072 | 22.74 | ***<0.001*** | 0.076 | 0.989 |
| Directional entropy | 1.47 | 0.253 | 15.91 | ***<0.001*** | 0.59 | 0.671 |
| Speed entropy | 5.77 | ***0.037*** | 5.34 | ***0.017*** | 1.26 | 0.303 |
| Entropy | 5.24 | ***0.045*** | 15.08 | ***<0.001*** | 0.14 | 0.967 |
| **Dark Cycle** | **Diet** | | **Body Part** | | **Interaction** | |
| **Parameter** | ***F*** | ***p*** | ***F*** | ***p*** | ***F*** | ***p*** |
| Center entries | 1.65 | 0.223 | 27.13 | ***<0.001*** | 0.10 | 0.983 |
| Center time (%) | 0.51 | 0.490 | 6.82 | ***0.014*** | 2.38 | 0.065 |
| Center distance (cm) | 2.81 | 0.120 | 42.37 | ***<0.001*** | 1.09 | 0.373 |
| Velocity (cm/s) | 2.49 | 0.141 | 191.51 | ***<0.001*** | 0.54 | 0.707 |
| Freezing time (%) | 1.57 | 0.234 | 14.63 | ***<0.001*** | 0.63 | 0.641 |
| Avoidance Index (ln) | 0.66 | 0.432 | 6.34 | ***0.015*** | 1.56 | 0.201 |
| Directional entropy | 2.28 | 0.157 | 10.76 | ***<0.001*** | 2.06 | 0.101 |
| Speed entropy | 2.21 | 0.163 | 114.44 | ***<0.001*** | 6.70 | ***<0.001*** |
| Entropy | 0.59 | 0.456 | 6.62 | ***<0.001*** | 1.93 | 0.120 |

**Supplemental Table 4. Female mixed-model ANOVA statistics for diet and body part movement**

| **Female Open Field Test Statistics (Diet x Body Part)** | | | | | | |
|---|---|---|---|---|---|---|
| **Light Cycle** | **Diet** | | **Body Part** | | **Interaction** | |
| **Parameter** | ***F*** | ***p*** | ***F*** | ***p*** | ***F*** | ***p*** |
| Center Entries | 0.90 | 0.366 | 69.58 | ***<0.001*** | 1.01 | 0.415 |
| Center Time (%) | 0.95 | 0.352 | 52.85 | ***<0.001*** | 1.19 | 0.330 |
| Center Distance (cm) | 0.09 | 0.774 | 77.32 | ***<0.001*** | 0.62 | 0.652 |
| Velocity (cm/s) | 1.79 | 0.210 | 197.66 | ***<0.001*** | 0.43 | 0.788 |
| Freezing Time (%) | 54.88 | ***<0.001*** | 66.25 | ***<0.001*** | 38.29 | ***<0.001*** |
| Avoidance Index (ln) | 1.31 | 0.279 | 87.10 | ***<0.001*** | 0.22 | 0.924 |
| Directional Entropy | 0.08 | 0.789 | 56.57 | ***<0.001*** | 0.42 | 0.790 |
| Speed Entropy | 9.71 | ***0.011*** | 3.82 | 0.059 | 0.88 | 0.484 |
| Entropy | 1.42 | 0.261 | 35.96 | ***<0.001*** | 0.06 | 0.993 |
| **Dark Cycle** | **Diet** | | **Body Part** | | **Interaction** | |
| **Parameter** | ***F*** | ***p*** | ***F*** | ***p*** | ***F*** | ***p*** |
| Center Entries | 3.88 | 0.077 | 25.01 | ***<0.001*** | 1.84 | 0.140 |
| Center Time (%) | 3.64 | 0.086 | 5.84 | ***0.022*** | 2.03 | 0.108 |
| Center Distance (cm) | 2.46 | 0.148 | 11.17 | ***0.003*** | 0.12 | 0.973 |
| Velocity (cm/s) | 1.75 | 0.216 | 68.97 | ***<0.001*** | 1.91 | 0.128 |
| Freezing Time (%) | 1.37 | 0.269 | 15.97 | ***<0.001*** | 1.03 | 0.405 |
| Avoidance Index (ln) | 3.57 | 0.088 | 6.86 | ***<0.001*** | 2.34 | 0.072 |
| Directional Entropy | 1.17 | 0.305 | 41.28 | ***<0.001*** | 1.55 | 0.207 |
| Speed Entropy | 1.35 | 0.272 | 66.82 | ***<0.001*** | 2.37 | 0.068 |
| Entropy | 3.66 | 0.085 | 6.37 | ***<0.001*** | 2.18 | 0.089 |

**Supplemental Table 5. Open field test behavioral PCA Loadings from mouse center**

| Open Field Test PCA Loadings (mouse center) | | | | | | |
|---|---|---|---|---|---|---|
| **Male** | **PC1** | **PC2** | **PC3** | **PC4** | **PC5** | **PC6** |
| Entry counts | 0.25 | 0.62 | 0.29 | 0.06 | 0.64 | 0.25 |
| Directional entropy | -0.35 | 0.40 | -0.53 | -0.22 | 0.24 | -0.57 |
| Avoidance Index (ln) | -0.33 | -0.61 | 0.08 | -0.19 | 0.68 | 0.06 |
| Freezing (%) | 0.61 | -0.26 | 0.07 | 0.29 | 0.20 | -0.65 |
| Velocity (cm/s) | -0.19 | 0.11 | 0.74 | -0.46 | -0.18 | -0.40 |
| Speed entropy | -0.54 | 0.08 | 0.26 | 0.78 | 0.01 | -0.15 |
| **Female** | **PC1** | **PC2** | **PC3** | **PC4** | **PC5** | **PC6** |
| Entry counts | 0.48 | -0.40 | 0.20 | 0.20 | -0.09 | 0.72 |
| Directional entropy | 0.42 | -0.15 | -0.57 | -0.61 | 0.31 | -0.01 |
| Avoidance Index (ln) | -0.54 | 0.20 | 0.17 | -0.42 | 0.33 | 0.59 |
| Freezing (%) | 0.37 | 0.54 | 0.12 | 0.35 | 0.66 | 0.00 |
| Velocity (cm/s) | -0.14 | -0.66 | 0.36 | 0.01 | 0.56 | -0.31 |
| Speed entropy | -0.36 | -0.20 | -0.68 | 0.54 | 0.21 | 0.20 |

**Supplemental Table 6. Mean (SEM) values for electrophysiology parameters**

| Electrophysiology Values Mean (SEM) | | | | |
|---|---|---|---|---|
| **Males** | **ZT 6-10** | | **ZT 18-22** | |
| **Parameter** | **CTR** | **aHFD** | **CTR** | **aHFD** |
| Rm | 139.43 (20.11) | 128.26 (11.65) | 102.54 (10.97) | 300.4 (20.71) |
| Cm | 60.14 (5.20) | 44.32 (5.60) | 79.79 (6.14) | 28.77 (1.40) |
| RMP | -66.87 (1.63) | -65.71 (2.50) | -66.87 (0.92) | -59.48 (1.56) |
| AP Threshold | -36.18 (1.76) | -36.7 (2.47) | -34.71 (1.10) | -39.32 (1.09) |
| Peak amplitude | 89.89 (2.50) | 89.67 (2.66) | 89.92 (3.21) | 90.53 (3.71) |
| Area | 317.66 (41.36) | 213.54 (65.32) | 434.65 (45.56) | 283.49 (65.00) |
| AHP | -1.57 (0.73) | -2.76 (1.08) | -1.02 (0.68) | -3.32 (1.32) |
| Half-width | 2.31 (0.18) | 2.01 (0.18) | 2.67 (0.16) | 2.52 (0.55) |
| Rise tau | 0.53 (0.04) | 0.48 (0.04) | 0.56 (0.03) | 0.61 (0.13) |
| Decay tau | 2.11 (0.15) | 1.98 (0.20) | 9.65 (7.02) | 2.51 (0.39) |
| Rheobase | 89.33 (14.69) | 100.00 (11.52) | 103.64 (17.07) | 48.57 (7.10) |
| G (-120 to -90mV) | 0.15 (0.02) | 0.16 (0.02) | 0.16 (0.03) | 0.12 (0.01) |
| G (0 to +30mV) | 1.65 (0.30) | 1.19 (0.13) | 0.78 (0.13) | 1.65 (0.16) |
| Reversal Potential | -80.57 (2.41) | -76.24 (5.95) | -73.32 (4.37) | -74.56 (3.77) |
| **Females** | **ZT 6-10** | | **ZT 18-22** | |
| **Parameter** | **CTR** | **aHFD** | **CTR** | **aHFD** |
| Rm | 160.00 (15.03) | 206.08 (10.12) | 207.44 (22.24) | 216.49 (18.73) |
| Cm | 39.31 (3.76) | 32.92 (3.06) | 38.68 (5.23) | 37.95 (5.95) |
| RMP | -62.849 (2.23) | -65.8 (1.42) | -69.45 (2.26) | -62.51 (1.95) |
| AP Threshold | -41.57 (2.80) | -38.84 (1.39) | -42.42 (1.80) | -39.35 (1.52) |
| Peak amplitude | 86.23 (3.40) | 93 (2.17) | 89.57 (3.96) | 89.53 (2.23) |
| Area | 108.8 (62.59) | 511.86 (32.16) | 367.07 (69.08) | 294.43 (48.60) |
| AHP | -6.94 (1.57) | 0.50 (0.88) | -3.08 (1.00) | -2.62 (0.99) |
| Half-width | 2.02 (0.25) | 2.27 (0.12) | 2.14 (0.15) | 2.30 (0.14) |
| Rise tau | 0.65 (0.10) | 0.87 (0.40) | 0.49 (0.03) | 0.50 (0.04) |
| Decay tau | 3.27 (1.27) | 2.18 (0.15) | 12.22 (10.34) | 2.42 (0.21) |
| Rheobase | 61.25 (10.64) | 69.03 (11.69) | 47.78 (14.12) | 79.41 (8.16) |
| G (-120 to -90mV) | 0.23 (0.03) | 0.17 (0.01) | 0.15 (0.01) | 0.17 (0.02) |
| G (0 to +30mV) | 1.81 (0.16) | 1.42 (0.15) | 1.47 (0.19) | 1.65 (0.16) |
| Reversal Potential | -71.06 (3.22) | -81.94 (2.33) | -83.88 (2.46) | -81.59 (2.43) |

**Supplemental Table 7. Statistics for electrophysiology parameters**

| Electrophysiology Statistics | | | | | | | | |
|---|---|---|---|---|---|---|---|---|
| **Males** | **N-Value** | | **Diet** | | **ZT** | | **Interaction** | |
| **Parameter** | **Mice** | **Cells** | ***F*** | ***p*** | ***F*** | ***p*** | ***F*** | ***p*** |
| Rm | 16 | 57 | 25.15 | ***<0.001*** | 16.30 | ***<0.001*** | 35.81 | ***<0.001*** |
| Cm | 16 | 57 | 44.75 | ***<0.001*** | 0.10 | 0.748 | 13.16 | ***0.001*** |
| RMP | 16 | 56 | 5.58 | ***0.022*** | 3.17 | 0.081 | 3.12 | 0.083 |
| Threshold | 16 | 52 | 2.42 | 0.126 | 0.09 | 0.768 | 1.51 | 0.225 |
| Amplitude | 16 | 52 | 0.00 | 0.949 | 0.02 | 0.890 | 0.02 | 0.895 |
| Area | 16 | 52 | 5.37 | ***0.025*** | 2.93 | 0.093 | 0.18 | 0.672 |
| AHP | 16 | 52 | 3.06 | 0.087 | 0.00 | 0.984 | 0.31 | 0.581 |
| Half-width | 16 | 52 | 0.44 | 0.509 | 1.69 | 0.200 | 0.05 | 0.829 |
| Rise tau | 16 | 52 | 0.00 | 0.984 | 0.97 | 0.329 | 0.42 | 0.522 |
| Decay tau | 16 | 52 | 1.26 | 0.267 | 1.64 | 0.207 | 1.15 | 0.289 |
| Rheobase | 16 | 51 | 2.74 | 0.105 | 1.90 | 0.175 | 6.28 | ***0.016*** |
| G (-120 to -90mV) | 14 | 52 | 0.53 | 0.470 | 1.08 | 0.304 | 1.61 | 0.210 |
| G (0 to +30mV) | 12 | 43 | 2.82 | 0.101 | 0.33 | 0.571 | 12.88 | ***<0.001*** |
| Reversal Potential | 14 | 52 | 0.12 | 0.731 | 1.30 | 0.259 | 0.47 | 0.495 |
| **Females** | **N-Value** | | **Diet** | | **ZT** | | **Interaction** | |
| **Parameter** | **Mice** | **Cells** | ***F*** | ***p*** | ***F*** | ***p*** | ***F*** | ***p*** |
| Rm | 17 | 75 | 3.45 | 0.068 | 2.77 | 0.100 | 1.29 | 0.259 |
| Cm | 17 | 75 | 0.76 | 0.386 | 0.31 | 0.582 | 0.39 | 0.535 |
| RMP | 17 | 75 | 0.47 | 0.494 | 0.33 | 0.567 | 6.27 | ***0.015*** |
| AP Threshold | 15 | 62 | 2.10 | 0.152 | 0.11 | 0.746 | 0.01 | 0.933 |
| Amplitude | 15 | 60 | 1.92 | 0.172 | 0.11 | 0.740 | 1.40 | 0.242 |
| Area | 15 | 60 | 15.85 | ***<0.001*** | 0.62 | 0.434 | 20.79 | ***<0.001*** |
| AHP | 15 | 60 | 15.01 | ***<0.001*** | 0.20 | 0.660 | 8.64 | ***0.005*** |
| Half-width | 15 | 60 | 1.49 | 0.227 | 0.14 | 0.708 | 0.05 | 0.819 |
| Rise tau | 15 | 60 | 0.22 | 0.643 | 1.17 | 0.285 | 0.14 | 0.708 |
| Decay tau | 15 | 60 | 2.20 | 0.144 | 1.25 | 0.268 | 1.82 | 0.183 |
| Rheobase | 15 | 62 | 2.29 | 0.135 | ***0.01*** | 0.917 | 1.05 | 0.310 |
| G (-120 to -90mV) | 17 | 75 | 1.67 | 0.201 | 3.65 | 0.060 | 3.17 | 0.079 |
| G (0 to +30mV) | 17 | 75 | 0.79 | 0.378 | 0.02 | 0.886 | 3.03 | 0.086 |
| Reversal Potential | 17 | 75 | 3.66 | 0.060 | 4.20 | ***0.044*** | 5.88 | ***0.018*** |

**Supplemental Table 8. PCA Loadings for electrophysiology basal and action potential properties**

| Electrophysiology PCA Loadings | | | | | |
|---|---|---|---|---|---|
| **Male** | **PC1** | **PC2** | **PC3** | **PC4** | **PC5** |
| Half-width | 0.47 | 0.14 | -0.09 | -0.10 | 0.10 |
| Rise tau | 0.46 | 0.19 | -0.05 | -0.13 | 0.26 |
| Area | 0.42 | -0.19 | 0.38 | -0.08 | -0.28 |
| AHP | 0.36 | -0.25 | 0.13 | -0.30 | -0.46 |
| AP Threshold | 0.25 | -0.04 | -0.44 | 0.49 | -0.33 |
| Decay tau | 0.17 | -0.03 | 0.46 | 0.44 | 0.47 |
| Cm | 0.07 | -0.44 | 0.25 | 0.38 | 0.09 |
| RMP | 0.06 | 0.49 | -0.10 | 0.44 | -0.26 |
| Rheobase | -0.07 | -0.44 | -0.40 | -0.16 | 0.25 |
| Rm | -0.12 | 0.45 | 0.32 | -0.27 | -0.02 |
| Amplitude | -0.38 | -0.13 | 0.29 | 0.12 | -0.39 |
| **Female** | **PC1** | **PC2** | **PC3** | **PC4** | **PC5** |
| Amplitude | 0.14 | -0.26 | -0.40 | -0.61 | -0.08 |
| Rise tau | 0.13 | 0.27 | 0.39 | -0.17 | 0.07 |
| Cm | 0.12 | 0.46 | -0.08 | -0.43 | 0.18 |
| RMP | 0.02 | -0.01 | -0.52 | 0.35 | 0.49 |
| Rm | -0.01 | -0.50 | 0.01 | 0.28 | -0.15 |
| Decay tau | -0.10 | -0.15 | 0.33 | -0.08 | 0.72 |
| Rheobase | -0.28 | 0.37 | -0.16 | 0.06 | -0.33 |
| Threshold | -0.35 | 0.21 | -0.44 | 0.00 | 0.22 |
| Half-width | -0.42 | 0.31 | 0.19 | 0.26 | -0.03 |
| Area | -0.51 | -0.24 | 0.21 | -0.29 | 0.07 |
| AHP | -0.55 | -0.21 | -0.08 | -0.22 | -0.09 |
